# Vertebrate features revealed in the rudimentary eye of the Pacific hagfish (*Eptatretus stoutii*)

**DOI:** 10.1101/2020.08.24.265124

**Authors:** Emily M. Dong, W. Ted Allison

## Abstract

Hagfish eyes are markedly basic compared to the eyes of other vertebrates, lacking a pigmented epithelium, a lens, and a retinal architecture built of three cell layers – the photoreceptors, interneurons & ganglion cells. Concomitant with hagfish belonging to the earliest-branching vertebrate group (the jawless Agnathans), this lack of derived characters has prompted competing interpretations that hagfish eyes represent either a transitional form in the early evolution of vertebrate vision, or a regression from a previously elaborate organ. Here we show the hagfish retina is not extensively degenerating during its ontogeny, but instead grows throughout life via a recognizable Pax6+ ciliary marginal zone. The retina has a distinct layer of photoreceptor cells that appear to homogeneously express a single opsin of the *rh1* rod opsin class. The epithelium that encompasses these photoreceptors is striking because it lacks the melanin pigment that is universally associated with animal vision; notwithstanding, we suggest this epithelium is a homolog of gnathosome Retinal Pigment Epithelium (RPE) based on its robust expression of RPE65 and its engulfment of photoreceptor outer segments. We infer that the hagfish retina is not entirely rudimentary in its wiring, despite lacking a morphologically distinct layer of interneurons: multiple populations of cells exist in the hagfish inner retina that differentially express markers of vertebrate retinal interneurons. Overall, these data clarify Agnathan retinal homologies, reveal characters that now appear to be ubiquitous across the eyes of vertebrates, and refine interpretations of early vertebrate visual system evolution.

**HIGHLIGHTS:**

- Hagfish eyes are *degenerate* but not *degenerating*, i.e. *rudimentary* but *growing*
- Retinal interneurons discovered implying ancestral hagfish had derived retinas & vision
- Despite lacking pigment, a Retinal Pigmented Epithelium homolog functions in hagfish
- Revised synapomorphies illuminate the dimly lit origins of vertebrate eye evolution

**GRAPHICAL ABSTRACT:** 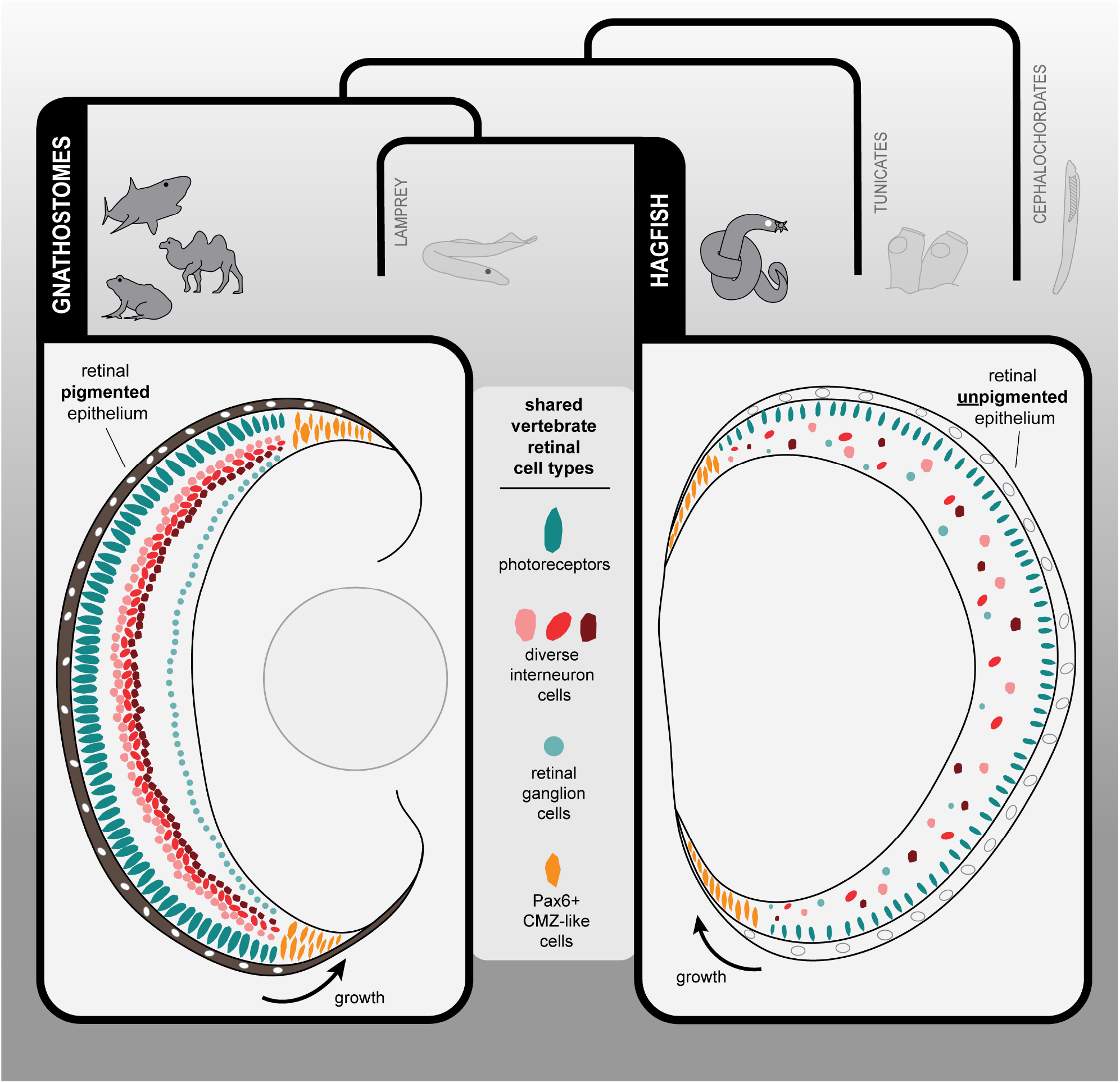

## INTRODUCTION

Vertebrate eyes are *remarkably* conserved and complex, appearing early in the evolution of this group as the familiar single-chambered camera-style organ with advanced optics at the anterior, and photoreceptors and retinal pigmented epithelium (RPE) lining the posterior of the eye’s globe. Across all vertebrate taxa, this morphology deviates little despite diverse visual ecologies and is easily recognizable even in early-branching vertebrates including the jawless lamprey. Homology of the eye across vertebrates is evident across various levels – anatomy, development, neural architecture, synaptic wiring, visual cell physiology, and molecular markers (Chow and Lang, 2001; Walls, 1942; Zuber, 2003). The eye is key to the success of vertebrates, yet its evolutionary origins have remained obscured.

Vertebrates are unique among chordates in sharing the following features (though in some derived instances features have been reduced or lost, e.g. in cavefish the eyes degenerate during ontogeny): a lens, extra-ocular musculature, and a neural retina supported by a pigmented retinal epithelium (RPE). The neural retina develops as an outpocketing of the forebrain and acquires key features that allow light to be translated into vision including a conserved neural architecture with three nuclear layers - the photoreceptors, interneurons and ganglion cells. Photoreceptors (rods & cones) transduce light into neural signals, interneurons (horizontal, amacrine & bipolar cells) compare these signals to compute and decant the various properties of the light stimuli, and retinal ganglion cells send the encoded information through their axons (the optic nerve) to the higher visual centres of the brain. A richly melanized RPE lines the back of the eye chamber, enveloping the photoreceptors, that functions to block stray photons and provide critical support for photoreceptor physiology (Strauss, 2005).

It is difficult to overstate (or briefly summarize) the impressive degree of conservation across vertebrate retinas at various levels of eye organization. While celebrated exceptions exist, such as rod-rich deep sea tube eyes (Collin et al. 1997) and four-eye fish (Borwein and Hollenberg, 1973), this vertebrate ocular *bauplan* is evident in lamprey, and thus has been retained since early vertebrate evolution prior to the advent of jaws or paired fins (limbs).

Hagfish are amongst the very rare exceptions to this canonical eye description in vertebrates and are particularly noteworthy because they populate an early branch on the vertebrate tree. Hagfish eyes, across two genera, are thus remarkable compared to all other vertebrates for the ocular characters they (seem to) lack, apparently concomitant with hagfish being positioned at the base of the vertebrate tree. Hagfish eyes are small, unpigmented and lack diagnostic jawed vertebrate features such as a duplex retina (with both rods & cones), lens, cornea and extraocular musculature (Locket and Jorgensen 1998, Lamb et al., 2007). The eye-cup is masked by semi-transparent or non-transparent overlying epidermis, the transparency of the skin varies across species (Fernholm and Holmberg, 1975). In some species, such as the Atlantic hagfish (*Myxine glutinosa*), the eye is also buried under body wall muscle (Fernholm, 1974). Though a lens and ocular musculature are absent, photoreceptors are present, as are retinal ganglion cells (Holmberg, 1971; 1970). Moreover, the hagfish neural retina lacks lamination into three nuclear layers, which is suggestive of a pineal-like architecture lacking interneurons, and of a simpler photoreceptive organ more suitable for detecting circadian light rhythms than for resolving images or vision.

The phylogenetic position and starkly rudimentary eye in hagfish have historically positioned them as a transitional form in early vertebrate visual system evolution; though an alternative view is that hagfish eyes may be rudimentary due to loss of characters. Delineating between these two explanations would fill a substantial gap in understanding how the vertebrate eye and its ‘inimitable contrivances’ evolved.

Here, we revisit the rudimentary hagfish retina to better appreciate the earliest steps in the history of vertebrate visual system evolution. Our objective was to explore retinal structure and identify cell types by molecular markers in the hagfish with a focus on Pacific hagfish (*Eptatretus stouii*) because its visual system seems somewhat less degenerate/rudimentary, compared to other species where the eyes are concealed by muscle (Fernholm and Holmberg, 1975; Jørgensen et al., 1998). Because hagfish embryos are prohibitively difficult to procure, we instead consider individuals across a range of sizes, including the smallest individuals attainable, to consider if hypotheses can be informed by characterizing these eyes over their ontogeny. We hypothesized that the hagfish eye possessed a greater number of conserved vertebrate eye characters than previously identified and that some (perhaps not all) features may be rudimentary as a result of secondary loss or arrested development. We reveal hagfish visual system characters that are otherwise conserved across vertebrate taxa, illuminating that some aspects of hagfishes’ remarkable eyes have regressed over evolutionary time.

## RESULTS

### The Pacific hagfish retina is simple and unpigmented

The eyes of Pacific hagfish (*E. stoutii*) are found beneath a layer of translucent skin (**Fig. 1A)**. They are small, embedded in the surrounding craniofacial muscle, and lack extra-ocular musculature attachments (**Fig. 1B,C**). Most notably, the eyes completely lack pigmentation **(Fig 1C,D)**. No lens is apparent, though the vitreous of the eye is protein-dense, evident by the high degree of hematoxylin staining (**Fig. 1D**). In accordance with an absent lens, no intraocular musculature is identifiable within the eye.

**Figure 1.**
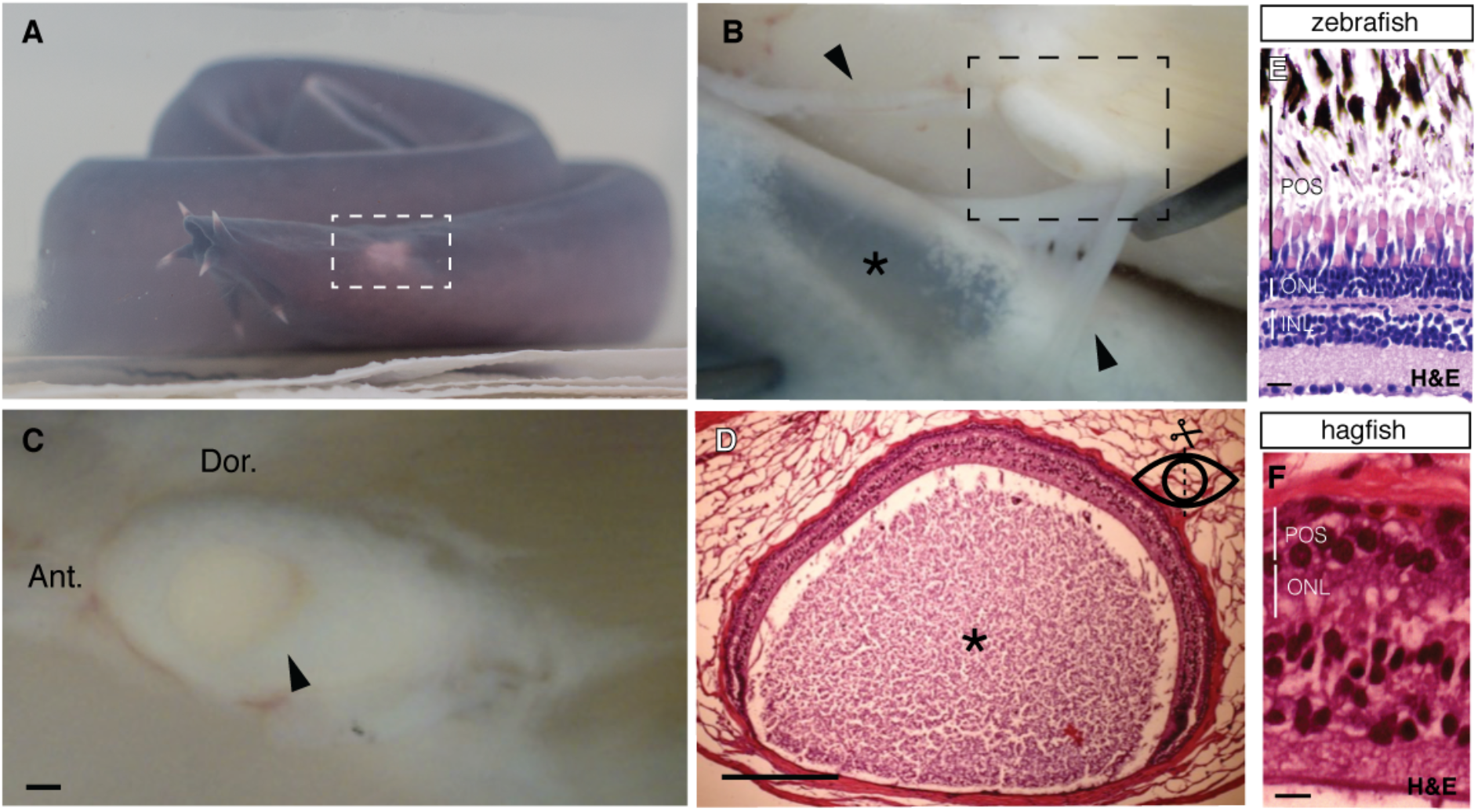
Eyes of Pacific hagfish are rudimentary, small, and strikingly devoid of melanin pigment. This sharply contrasts the eyes of all other vertebrate taxa and is concomitant with the position of hagfish as the earliest branch on the phylogeny of extant vertebrates. **(A)** The unpigmented eyes of Pacific hagfish are contrasted against a richly pigmented body epidermis. The epidermis is reduced to thin transparent windows of tissue covering each eye (boxed area; see also Supplemental Fig. S1). **(B)**. As revealed during dissection, the transparent window of epidermis (*) is held in position over each eye (boxed area) by connective tissue (arrowheads), and otherwise the epidermis is loosely attached to the body throughout most of the animal’s length. **(C)** The diminutive eye is embedded amongst the surrounding head musculature and is not mobilized by any extraocular muscles. Arrowhead indicates eyecup aperture (‘pupil’). Panels B & C dissect specimens fixed in paraformaldehyde, Fig. S1 shows similar with fresh tissue. **(D)** Eyecup of Pacific hagfish is rudimentary, surrounded by body wall muscle and collagen. Intra-ocular musculature is absent. A crystalline lens is not observed, though a protein rich (hematoxylin stained) accessory structure is present (*), perhaps analogous to vitreous or lens of gnathostomes. **(E, F)** Retina of Pacific hagfish lacks both pigment and the distinct three-layer neural architecture (photoreceptors, interneurons, ganglion cells) that are obvious and conserved in most all vertebrate retinas (exemplified here by zebrafish). POS, photoreceptor outer segment layer; ONL, outer nuclear layer; INL, inner nuclear layer. Scale bars are in C-D are 0.25mm, E-F are 10μm

In cross section, the typical vertebrate neural retina (exemplified here by zebrafish) is stratified into three distinct cellular layers (ONL, INL GCL) divided by two synaptic layers (IPL, OPL) (**Fig 1E**). In contrast, the retina of *E. stoutii* is not morphologically separated into this same layered organization (**Fig 1F**). While the precise identities of cells across the width of the retina are a subject of the current investigation, photoreceptors are found at the scleral-most region, while presumptive retinal ganglion cells are found basal to this, towards the vitreous (**Fig. 1F)**. Overall, the Pacific hagfish eye is rudimentary compared to exemplar eyes of other vertebrate taxa (see Introduction), consistent with past reports in various hagfish species (Fernholm and Holmberg, 1975; Lamb et al., 2007; Locket and Jørgensen, 1998).

### Hagfish eyes grow over ontogeny perhaps via Pax6-expressing progenitors at the retinal margins

To characterize if the rudimentary eyes of hagfish are changing over ontogeny, perhaps akin to degenerating eyes of other vertebrates adapted to dim-light environs (e.g. cavefish in which the eye diminishes significantly), we measured eye size in Pacific hagfish caught off the West coast of Canada. Because the hatching and raising of hagfish embryos for experimental purposes is rarely attainable, we collected wild-caught individuals and used fish size as a proxy for age. The eye was found to increase in both length and mass during ontogeny, and over a broad range of fish sizes **(Fig. 2F-G**; e.g. >4-fold increase in fish mass or length). Thus, the Pacific hagfish eyes appear degenerate (rudimentary and diminutive) but are not overtly degenerating over ontogeny.

**Figure 2.**
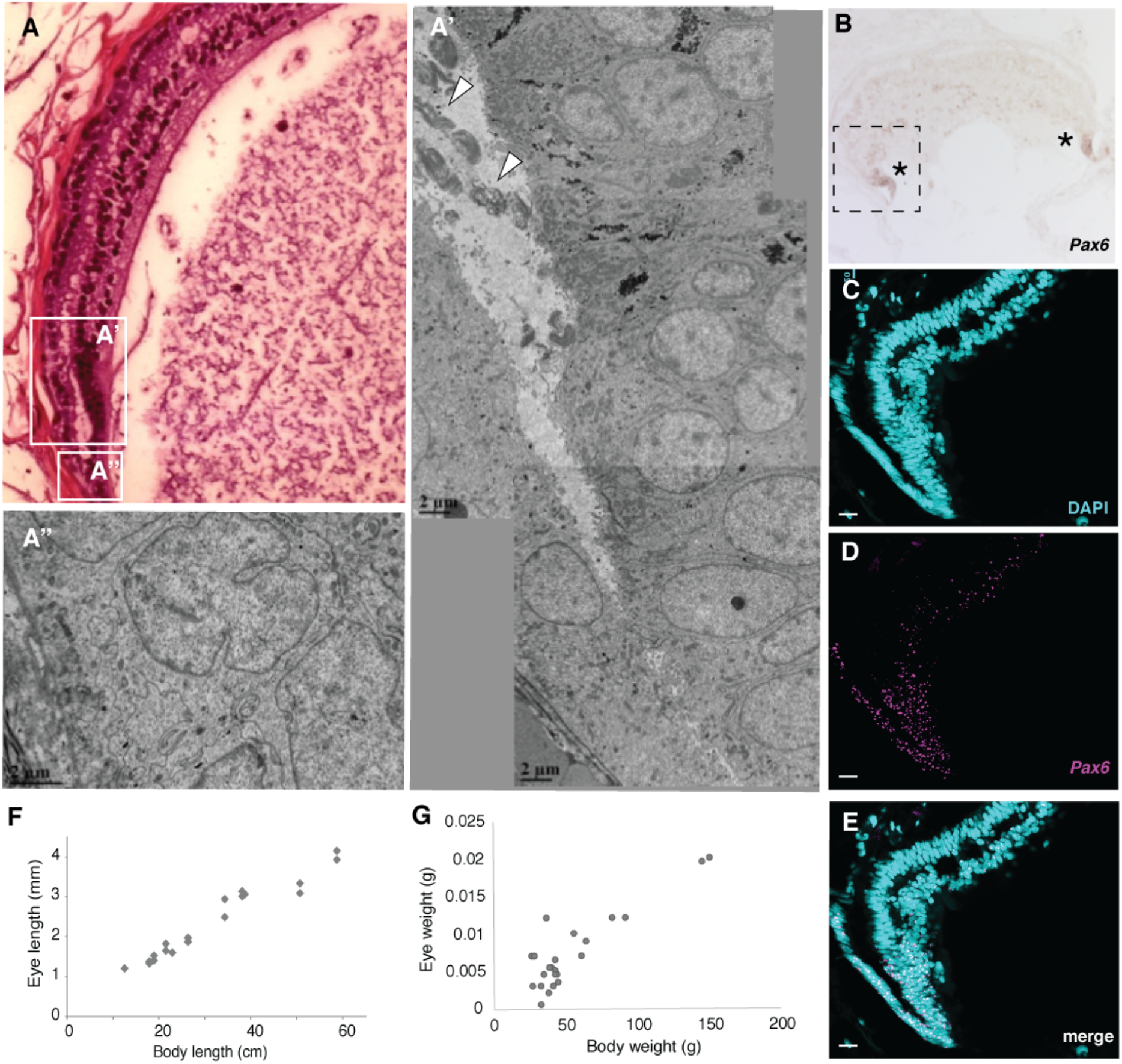
Hagfish retina has a peripheral tissue expressing *PAX6* that presumably contributes to eye growth late into ontogeny. This tissue is highly reminiscent of the multipotent and proliferative Ciliary Marginal Zone (CMZ) that supports eye growth late into ontogeny of early-branching gnathostomes. **(A)** Peripheral hagfish retina, with laminated central retina (at top) merging into a simpler neuroepithelium near the pupil (bottom). The cells located more peripherally appear less differentiated and organized, while those located more centrally are seen within distinct photoreceptor outer segments (indicated by arrowheads), and begin to laminate into discernable cell layers (A”). **(B)** Expression of *PAX6* transcript is highly enriched at the periphery of the eyecup (purple precipitate at both extreme tips of this sectioned retina, *). **(C-E)** *PAX6* transcript (magenta) is expressed by the majority of cells at the peripheral margin of the CMZ. View is equivalent to the boxed region in panel B, nuclei are counterstained with DAPI (cyan), scale bars are 100μm. **(F, G)** Pacific hagfish eyes grow throughout ontogeny. Each dot represents one eye from one individual animal, and two different groups of animals are presented in panels F vs. G. A nearly isometric increase in eye size is observed over a >4-fold increase in animal length or mass.

Observing larger eyes in larger hagfish was reminiscent of early-branching gnathostomes where eye growth continues late into the animals’ ontogeny, supported by a population of proliferating undifferentiated cells at the outer edges of the mature retina, known as the ciliary marginal zone (CMZ) (Fischer and Reh, 2000; Kubota et al., 2002; Sánchez-Farías and Candal, 2016; Sukeena et al., 2016; Villar-Cheda et al., 2008; Wetts et al., 1989). We observed that hagfish eyes possess a comparable region and propose that it contributes to the sustained growth of the eye throughout the lifetime of the fish. In hagfish, the cells at the retinal margins lack photoreceptor outer segments and have irregularly shaped nuclei distinct from other retinal cells in the central areas of the retina (**Fig. 2A**). A gradient of more elaborate cell morphologies is observed closer to the center of the retina, with increasing resemblance to fully differentiated cells; this gradient is reminiscent of observing more mature central retina in teleost fish (**Fig. 2A’-A’’**). The simpler retinal neuroepithelium at the margins is observed to merge with a more distal (scleral) epithelium (the presumptive RPE, described below). This merged retinal margin is positioned at the edge of the eye cup, comparable to the gnathostome CMZ where it would be contiguous with tissue forming the pupil.

In gnathostomes, the CMZ expresses *PAX6* (Fischer et al., 2014). An ortholog of *PAX6* highly conserved with that of other vertebrates, was found to be expressed in the hagfish retina. *PAX6* expression is most concentrated at the tip of the retinal marginal zone (**Fig. 2B-E**), suggesting that not only do these cells appear undifferentiated by morphology, but that they also express transcripts related to the maintenance of a multipotent cell state (Fischer et al., 2014; Marquardt et al., 2001).

### Interneurons are found despite the lack of vertebrate-typical retinal layering

Apart from photoreceptors and retinal ganglion cells, the cell identities within the hagfish retina have remained obscured and a segregated interneuron layer in hagfish is not obvious. This simple retinal composition noted in various hagfish species is interpreted by some to reflect a primitive neural architecture more reminiscent of a the two-layered morphology and physiology of the pineal organ (Lamb et al., 2007). We observed that the inner retinal cells below the photoreceptor layer in Pacific hagfish are a diverse mix of cells that possess nuclei that are not homogenous in shape or size, with diverse heterochromatin patterning (**Supp. Fig 2**). We hypothesized that additional cell types aside from retinal ganglion cells populate this region. RNA-sequencing indicated expression of putative interneuron cell markers (**Fig.3A**). Distribution of these transcripts was assessed via *in situ* hybridization on sectioned hagfish retina (**STAR Methods**). Expression of *PKC-α* (**Fig. 3E**, a marker of vertebrate bipolar cells), *PAX6* (**Fig. 3B**, a marker of vertebrate amacrine cells and retinal ganglion cells) and *CALBINDIN* (**Fig. 3F**, a marker of vertebrate horizontal cells) were each found to be expressed in a subset of cells in the inner nuclear layer. Each of these genes are high-fidelity markers of retinal cell types across diverse vertebrates (Greferath et al., 1990; Marquardt and Gruss, 2002)

As a separate challenge to the hypothesis that hagfish retinas have a simple neural architecture and lack interneurons, we characterized the distribution of the synapses. We observed immunoreactivity for synapse marker SV2 (synaptic vesicle protein 2) to localize into two layers of hagfish retina (**Fig. 3C**, **Supp. Fig 3**), one being associated with the basal end of photoreceptors, and a second layer intermingled with nuclei closer to the vitreous. Staining of F-actin (enriched in synapses across diverse animals) using fluorescently-labelled-phalloidin also supported the existence of multiple synaptic layers (**Supp. Fig 3**). Though less precisely organized, this synaptic architecture is reminiscent of jawed vertebrate retinas, where two distinct plexiform layers demarcate the presence of interneurons connecting the photoreceptor and ganglion cells.

**Figure 3.**
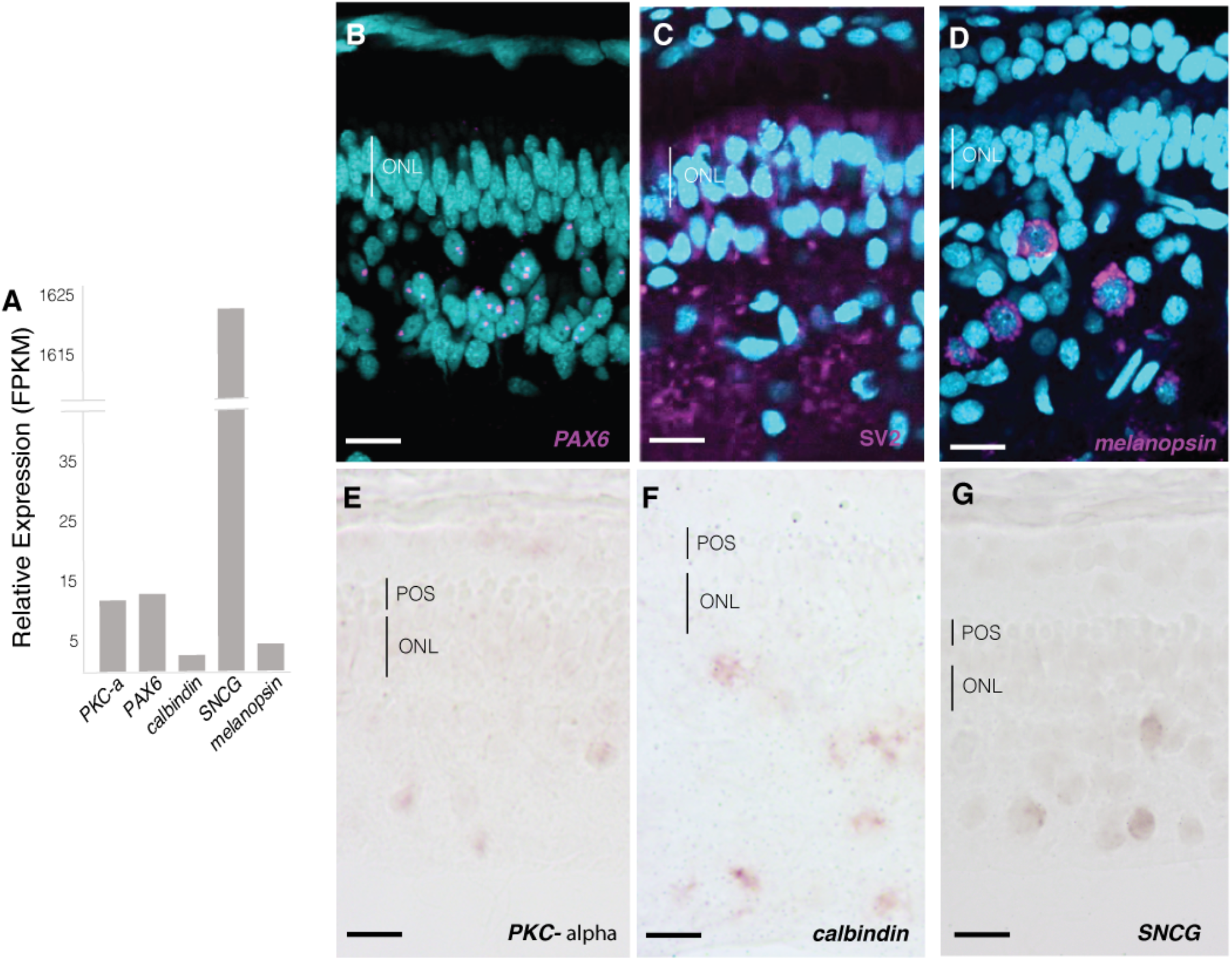
A subset of cells in the Pacific hagfish retina each express markers of various gnathostome retinal interneurons or retinal ganglion cells. Within the inner retinal layer (positioned closest to the vitreous), a variety of cell morphologies are observable (Fig S2) and a small proportion of cells each express genes that reliably serve as markers of interneuron subtypes (bipolar, amacrine & horizontal cells – Supplemental Table 1) across diverse vertebrates. **(A)** Expression of classic retinal interneuron markers detected in hagfish eyes by RNA-Seq. **(B)** Transcripts of *PAX6* are observed within the nuclei of a subset of inner retina cells, suggestive of amacrine cells. **(C)** The identification of interneurons in retina of Pacific hagfish is supported by the detection of synaptic markers (immunoreactivity for SV2, see also Supplemental Figure S3). **(D)** Detection of melanopsin transcript surrounding the nucleus in a subset of inner retina cells implies presence of ipRGCs (intrinsically photosensitive Retinal Ganglion Cells). **(EG)** Expression of transcripts encoding *PKCα, Calbindin*, or *γ-synuclein* (*SNCG*) each in a subset of inner retina cells is consistent with their use as retinal cell markers across diverse vertebrates, where they identify bipolar cells, horizontal and ganglion cells, or ganglion cells, respectively. DAPI nuclear stain is in cyan. Scale bars are 20μm.

Retinal Ganglion Cells (RGCs) have previously been examined in the hagfish retina (Holmberg, 1971; Kobayashi, 1964; Sun et al., 2014). We characterized the distribution of RGCs by in situ hybridization for cell-type specific marker *γ-synuclein (SNCG*, **Fig. 3G**) (Laboissonniere et al., 2019; Soto et al., 2008) and *melanopsin* (**Fig. 3D**). RGCs represented a subset of cells identified within the inner retina cell layer, distributed both at the vitreal retinal margin and in more apical locations within the inner retina cell layer.

In sum, the inner layer of hagfish retina contains cells of various types, including subsets of cells expressing markers of retinal ganglion cells, amacrine cells, bipolar cells and horizontal cells. These cells intermingle with immunoreactivity for synaptic markers that localize in a layer distinct from the photoreceptor synapses. Previous characterizations have described hagfish retinas as having only two layers of cells and speculated that this simplicity is comparable to the vertebrate pineal; the data presented here reveal the hagfish retina has greater complexity, and recognizable neuronal features conserved with retinas of jawed vertebrates.

### A homolog of RPE, the RnPE, interdigitates with photoreceptors lining the hagfish outer retina

In jawed vertebrates, the retina is sustained by the retinal pigment epithelium (RPE), a monostratified epithelium with dense melanin, located between the photoreceptor outer segments and the choroid (a network of vasculature that supports the outer retina). This vital partnership between the neural retina and RPE is unique to vertebrates.

We sought to determine if hagfish photoreceptors, which appear morphologically rod-like (**Fig. 4E**), express the rod associated opsin—rhodopsin. Rhodopsin (RH1) is the only visual opsin known to be present in hagfish eyes, though its expression pattern and the presence of other visual opsins in the genome remain to be determined (Lamb et al., 2016). Through RNA sequencing we identified only one visual ciliary opsin: a highly expressed *RH1*. Homology searching through the *de novo* assembled transcriptome using HMMER (Eddy, 2011) and BLASTP 2.6.0 (Altschul et al., 1997) did not reveal any additional visual opsins. In comparison with other vertebrates, *E. stoutii RH1* shares high sequence similarity, and is clustered among other vertebrate *RH1s* with high certainty (**Supp. Fig. 4**).

**Figure 4.**
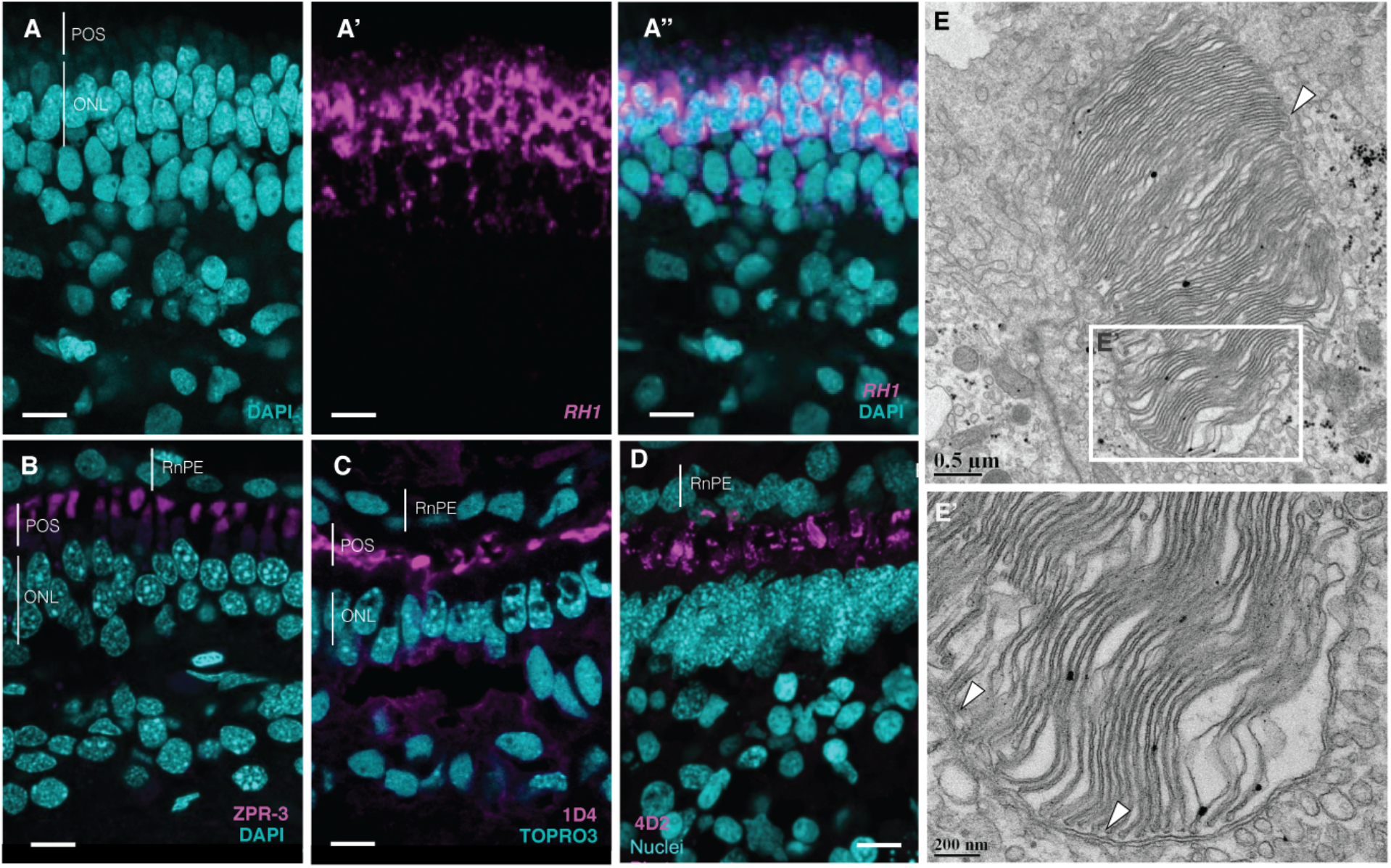
Photoreceptors in the outer retina of Pacific hagfish express rod opsin transcript and immunoreactivity. The expression of *RH1* rod opsin appears homogeneous, i.e. no cells in this layer are observed to lack expression. **(A)** *RH1* transcript (magenta) is robustly expressed in a layer of outer retina cells. **(B-D)** Immunoreactivity for *RH1* protein (magenta), detected with any of three antibodies against gnathostome rod opsin (ZPR-3, 1D4, 4D2, respectively) label a cell compartment (the outer segment) positioned apical of the transcript, suggesting polarized ciliary photoreceptor morphology similar to gnathostome rods and cones. Nuclei are counterstained with DAPI or TP-PRO-3 (cyan). Scale bars are 20μm. **(E) Ultrastructure of** photoreceptor show outer segments with membranous discs similar to rod, where a surrounding membrane exterior to the discs is indicated by arrows. POS, photoreceptor outer segment layer; ONL, outer nuclear layer; RnPE, retinal non-pigmented epithelium.

*In situ* hybridization of hagfish *RH1* using RNA probes designed from transcriptome derived sequences showed highly specific staining in the photoreceptors in close association with the nuclei (**Fig. 4A-C**). *RH1* appears to be expressed in all photoreceptors, making the presence of other ciliary photoreceptor types unlikely. Additionally, the photoreceptor outer segments are readily immunolabelled by ZPR-3 (zebrafish rod opsin antibody) and 1D4 (monoclonal antibody raised against bovine rhodopsin C-terminus) and 4D2 (monoclonal antibody raised against bovine RH1 N-terminus) (**STAR methods, Fig. 4 D-E**). The protein exhibiting rod-opsin-immunoreactivity was localized to cellular compartments located distally (scleral side) of the *RH1* transcript localization, i.e. in the photoreceptor outer segments. This location was also noted as being immediately apical of an actin-rich retinal strip (**Supp. Fig. 3D**) that is positioned identically to the outer limiting membrane (composed of Müller glia end feet and their actin-rich cell adhesion with photoreceptors) in gnathostomes. Overall this polarized cell organization is exactly as is seen in the highly derived and polarized gnathostome rod (and cone) photoreceptor cells.

A single row of epithelial cells at the distal/outer extreme of the hagfish retina, beyond the photoreceptors has been unsatisfactorily characterized. Previous studies have shown that these cells possess microvilli on their basal surface in close association with nearby photoreceptor outer segments (Fernholm and Holmberg, 1975; Holmberg, 1971). This tissue has a similar location and organization to the retinal pigment epithelium (RPE) found in other vertebrates, but is strikingly disparate (even macroscopically, **Fig. 1, S1**) due to its complete lack of melanin pigment. Melanin pigment found in the RPE of vertebrates is key to sustaining healthy function in the eye. Curiously, though the eye is unpigmented, we have found via RNA-Seq that pigment-related genes are expressed in the eye, including *tyrosinase* (necessary for melanin synthesis) and *PMEL* (pre-melanosome protein necessary for stable melanin deposition in the melanosome) (**Fig. 5A**). Thus, we have termed this non-pigmented RPE equivalent the “Retinal (non)Pigment Epithelium” or RnPE.

**Figure 5.**
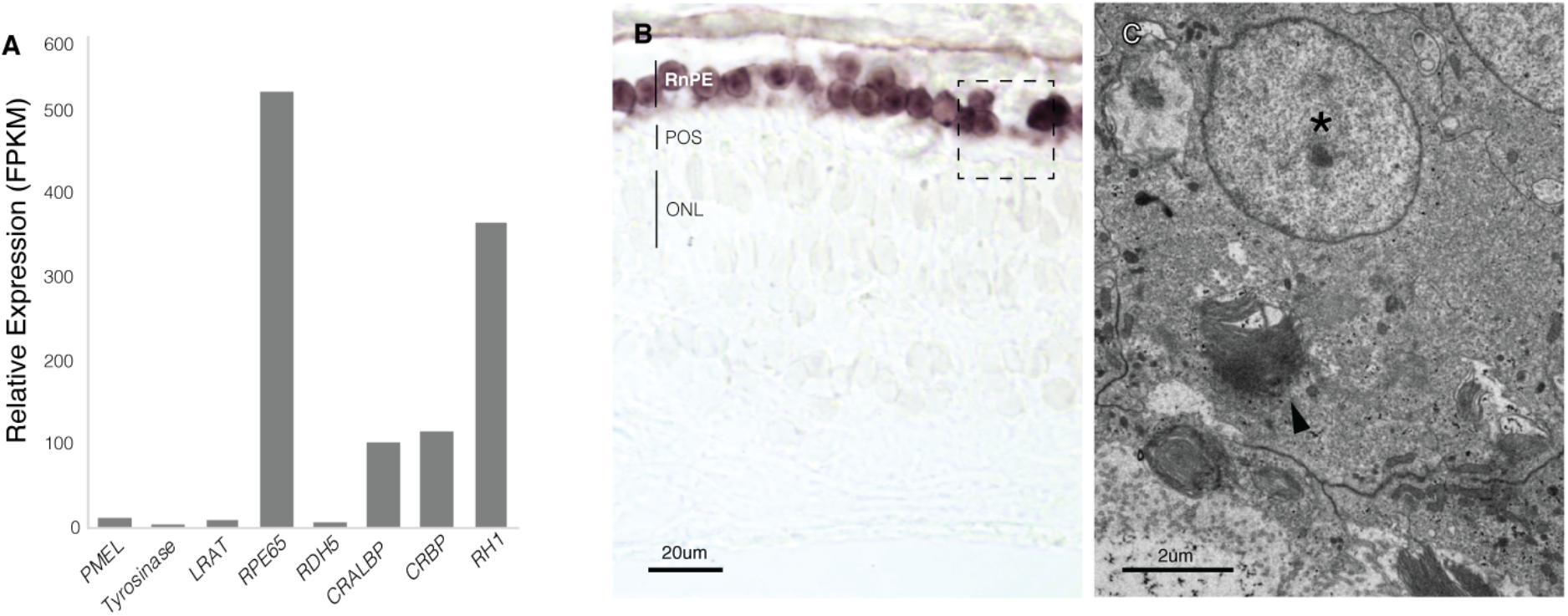
The outermost aspect of Pacific hagfish retina is encapsulated by an equivalent of gnathostome Retinal Pigment Epithelium (RPE) that starkly lacks pigment. We denote this as the RnPE, the Retinal non-Pigmented Epithelium. **(A)** Various eye transcripts detected by RNA-Seq are suggestive of RPE physiology and vestigial expression of machinery to produce melanin pigment. **(B)** Robust expression of *RPE65* transcript, a marker of RPE cells, is detected in the outer retinal epithelium (hagfish RnPE), positioned on the scleral side of photoreceptor outer segments (POS). **(C)** Phagocytosis of expired portions of the outer segments of the photoreceptors is seen in the hagfish RnPE. A portion of an outer segment has engulfed by a phagosome is indicated by the arrowhead, an asterisk denotes retinal epithelium nucleus.

A major function of the RPE is to maintain the integrity of the photoreceptor outer segments by phagocytosing shed photoreceptor discs. This process renews the photoreceptors and in part prevents oxidative stress. As previously documented (Holmberg, 1971), we have found ultrastructural evidence of phagosome-like organelles in the hagfish RnPE, which contain what appears to be pieces of photoreceptor outer segments (**Fig. 5C**).

The RPE also serves a highly necessary role in retinoid cycling – specifically, the production of 11-cis retinal, a chromophore associated with visual opsins that is required for visual function. Phototransduction cannot be sustained without the renewal of 11-cis retinal by the RPE. Retinoid cycling is achieved by the enzymatic action of lecithin-retinol acyltransferase (LRAT), 11-*cis*-retinol dehydrogenase 5 (RDH5) and *RPE65*, supported by cellular retinaldehyde-binding protein (CRALBP) (Batten et al., 2004; Moiseyev et al., 2003; Strauss, 2005). The transport proteins cellular retinol binding protein (*CRBP*) and interphotoreceptor retinoid-binding protein (IRBP) are also involved in the transport of highly hydrophobic retinoids between the RPE and photoreceptors (Strauss, 2005). Analysis of our transcriptome revealed that homologs of *RPE65, LRAT, RDH5, CRALBP* and *CRBP* are expressed in the hagfish eye (**Fig. 5A**). *IRBP* could not be identified. Further, we found that the expression of *RPE65* is highly specific and robustly expressed within all cells of the RnPE (**Fig. 5B**). In sum, the presence of phagosomes, the interdigitiation with polarized ciliary photoreceptors, the presence of melanin synthesis machinery, and the localized expression of conserved markers of RPE cells support the RnPE being a homolog of gnathostome RPE that was pigmented in the last common ancestor of hagfish and jawed vertebrates.

## DISCUSSION

### Illuminating conserved features that Hagfish eyes share with other vertebrates

The hagfish eye rudiment has remained largely enigmatic, limiting an appreciation for the early history of vertebrate eye evolution. Though apparently rudimentary in comparison to the elaborate eyes that are conserved across vertebrate taxa, hagfish eyes appear advanced when compared to non-vertebrate chordates, i.e. the photoreceptor clusters of tunicates and cephalochordates (though even these early-branching chordates show the conserved requirement for pigment in close association with photoreceptor cells). This graduating pattern of character complexity is to some degree consistent with aspects of hagfish eyes representing a transitional form in visual system evolution. However, the data here strongly support that some species of extant hagfishes have eye characters that are homologous with those of jawed vertebrates, and therefore were present in their last common ancestor. Thus, it appears that several ocular characters have regressed in hagfish, rendering them difficult to recognize despite decades of interest.

The data here reveal several cell- and tissue-level characters in the hagfish retina that are familiar from the eyes of fishes, birds and mammals. At the level of tissues, we have gained an appreciation that the hagfish retina has a Pax6+ growth zone that is retained late into ontogeny, as is observed robustly in other adult fishes and in developing amniotes. The RnPE is present and functional, despite the hagfish eye being most striking due to the stark absence of melanin pigment. Beyond the retina, several conserved ocular tissues remain enigmatic in hagfish: Homologs of ocular muscle are not apparent, and instead of a crystalline lens a less impressive soft tissue is present (we cannot yet rule out that it might play some role in collecting, filtering and/or focusing incoming photons).

At the cellular level, homologs of vertebrate retinal ganglion cells (and optic nerve) had been recognized previously in hagfish and had been clarified to project to visual centres instead of (or perhaps in addition to) projecting to pacemaker nuclei reserved for circadian rhythms (Wicht & Northcutt, 1990). On the other hand, more recent work revealed that some hagfish retinal cells express melanopsin and are likely equivalent to ipRGCs and support the hagfish retina as having photoreceptive roles in circadian rhythm. Otherwise, identifying cell types in Hagfish retina has been limited to photoreceptors, characterized by ultrastructure or more rarely by immunostaining.

The apparent simplicity of the hagfish retina stands in contrast to the three nuclear cell layers that are ubiquitous across vertebrate retina. Typical vertebrate retinas share an outermost layer comprised of photoreceptors, middle layer of interneurons (horizontal, bipolar, and amacrine cells), and an innermost layer containing retinal ganglion cells. In order for images to form out of the chaos of visual information hitting the retina, neural computations occur via the highly complex relationships that exist among photoreceptors and interneurons. Hagfish retinas lack these three morphologically distinct cellular layers and it is this obscured retinal lamination that has likewise obscured the presence of an interneuron cell population (Lamb et al. 2007), relegating the hagfish retina to a limited photoreceptive ability and precluding any history of advanced vision. However, the data here reveal that hagfish possess retinal interneurons by identifying the presence of *CALBINDIN, PAX6* and *PKC-α* labelling each within a subset of cells in the inner nuclear layer. Moreover, synaptic markers localized into two well-demarcated retinal plexiform layers independently supports the presence of retinal interneurons. Overall, at least one species of extant hagfish has a neural architecture in its retina that is much more elaborate than previously recognized and characterized by exact parallels with the retina features conserved across jawed vertebrates. It is less parsimonious to speculate these striking similarities arose via convergent evolution, and so the last common ancestor of hagfish, lampreys and gnathostomes most likely had a retina with neural architecture (and concomitant visual ability) similar to modern jawed vertebrates. At least in this aspect, the hagfish retina appears to have regressed over evolutionary time (perhaps associated with its deep-sea dim light habitats).

Hagfish photoreceptors appear simpler than those of other vertebrates, found either as a whorled structure in *Myxine* species, or shortened and non-tapered in *Eptatretus* species (Holmberg, 1971). The unique morphological qualities of these photoreceptors have obstructed their identification in the context of rod and cone identity. We have shown that the ultrastructurally rod-like photoreceptors of *E. stoutii* express rhodopsin transcript, further supporting them as rod-like. Moreover, robust outer segment immunoreactivity by 1D4, 4D2 (bovine rhodopsin) and ZPR-3 (zebrafish rod-labelling) antibodies suggests that these rhodopsin transcripts are translated and trafficked to the outer membranes, where they are well-positioned to elicit phototransduction. Rhodopsin (RH1) or rod opsin, is the only identified visual opsin in hagfish, and has been proposed to act through a phototransduction cascade similar to that of other vertebrates (Lamb et al., 2016; Vigh-Teichmann et al., 1984). Our work elevates and expands upon studies characterizing hagfish photoreceptors as rods by their rod-typical spectral sensitivity, and electrophysiological response to low light stimuli (Kobayashi, 1964; Steven, 1955).

The RPE provides a number of services to the photoreceptors including retinoid cycling, nutrient and metabolite exchange between vasculature and the retina, and phagocytosis of sloughed photoreceptor outer segments. These two interdigitated tissues co-exist in a highly intertwined relationship which, when disrupted, results in malfunction and disease (Strauss, 2005). While the presence of rod-like photoreceptors in hagfish suggests the retina can receive and transduce light signals, our identification of the presence of a retinal (non)pigment epithelium further implies that the continued function of photoreceptors can be sustained. The presence of ciliary opsin expressing photoreceptors in conjunction with an epithelial cell layer capable of retinoid cycling is a derived feature of chordate eye evolution, and exclusive to vertebrates. This partnership is absent in early diverging chordates such as tunicates, where visual cycle components are expressed within the photoreceptor cells (Tsuda et al., 2003). The shared presence of retinoid cycling components within a separate epithelial tissue in hagfish, lamprey and gnathostomes suggests that this RPE-photoreceptor partnership was present in their last common ancestor and predates the divergence of jawed from jawless vertebrates.

### Degenerate eyes that are not degenerating – ocular growth late into ontogeny

Post-natal eye growth has been lost in crown group vertebrates, but ongoing adult retinal neurogenesis is a broadly conserved and prominent feature in fishes. Across hagfish species a gradient of eye regression is apparent, with Atlantic species (*Myxine*) exhibiting small eyes buried under muscle and often unobservable when examining intact animals (field guides supporting species identification are thus not entirely incorrect when they say that hagfish lack eyes), and *Eptatretus* species which show far fewer restrictions in receiving environmental light {Fernholm:1975ct}. Pacific hagfish, in contrast to reasonable expectations of eye degeneration over ontogeny, were found to exhibit macroscopic growth of the eye isometric with body growth. Our data suggest that this growth is supported by a multipotent cell niche at the retinal margin, the ciliary marginal zone (CMZ). The cells at the CMZ of other early-branching vertebrates are capable of proliferating and adding to the fully formed retina post-development (Fischer and Reh, 2000; Hollyfield, 1971; Johns and Easter, 1977; Kubota et al., 2002). As seen in other vertebrates, the cells at the ciliary marginal zone express *PAX6* (a marker of undifferentiated multipotent progenitor cells), (Marquardt et al., 2001).

The CMZ in other vertebrates, viewed as a snapshot in time via retinal histology, can provide a proxy timeline for developmental events beginning peripherally as proliferating cells produce new neuroepithelium. Toward the central mature retina this gives rise to the retinal layers – RPE and neural retina, with the latter being composed of multiple glial and neuronal cell types decanted into three layers. Studying hagfish CMZ could circumvent several challenges of hagfish evo-devo. Considering that hagfish embryos are infamously difficult to attain, further work is warranted in exploring the hagfish CMZ as a window into retinal development. The data presented here regarding hagfish CMZ already suggest a conserved developmental source and common evolutionary ancestry for key retinal tissues (RPE, photoreceptors, retinal lamination) across the gnathostomes.

### Growing support for loss of missing eye characters

The absence of key ocular characters in the hagfishes, revealed here to be the result of character loss and eye regression, can now be rationalized against the phylogeny of agnathans. The ill-defined taxonomic relationships of agnathans have contributed much to the perplexity regarding the rudimentary hagfish eye. Until recently, it was debated whether hagfish and lamprey were a monophyletic group, or independent branches off the base of the vertebrate tree. Monophyly of hagfishes and lampreys is now strongly supported (Delarbre et al., 2002; Heimberg et al., 2010; Kuratani et al., 2018; Miyashita et al., 2019; Ota et al., 2011; 2007; Takezaki et al., 2003; Yu et al., 2008). Our data reinforce this conclusion, clarifying that vertebrates share key visual system characters (RPE, retinal architecture, eye growth) though some of these synapomorphies were concealed by evolutionary regression in the hagfish lineage. Moreover, monophyly of the agnathans greatly weakens the hypothesis that hagfish represent an ancient transitionary eye, because this interpretation requires that lamprey acquired vertebrate-like ocular characters independently. It is instead more likely that the last common ancestor of jawed vertebrates, lamprey and hagfishes all shared these derived characters and the hagfish eye is an extraordinary example of eye regression.

Widespread expression of *RH1* in *E. stoutii* photoreceptors is a key support in the hypothesized regression of hagfish eye features. Typical vertebrate photoreceptors express any one of five visual opsins, with rods expressing rhodopsin (*RH1*) and cones express a number of opsins covering the visual light spectrum (*RH2, LWS, SWS1, SWS2*) (Bowmaker, 2008). Each opsin is tuned to a particular wavelength allowing for colour and high acuity daylight vision in the case of cones, and dim light in the case of rods. Phylogenetic analysis supports rhodopsin as the most recently evolved visual opsin, arising from a duplication of RH2 (Okano et al., 1992). The lone presence of *RH1* expressing photoreceptors in hagfish demonstrates a lost capacity to express other visual opsins, given that their photoreceptor complement is likely to have more closely resembled lamprey, in which some species which possess all five visual opsins (*RH1, RH2, LWS, SWS1, SWS2*) (Collin et al., 2003). Future work should strive to identify lingering cone networks, if any exist. Leveraging genomic studies will be particularly necessary in interrogating the full extent of opsin gene loss.

The absence of pigment is one of the most striking features in the hagfish eye. Fossil data for hagfish suggests that the eyes of at least one basal hagfish species possessed pigment implying a pattern of regression in extant species (Gabbott et al., 2016). Oddly, though melanin is lost from the RPE, the machinery for producing melanin-rich tissues is obviously intact and functional in hagfish skin (except overlying the eye). Further, we have shown that the retina expresses pigment related transcripts including *PMEL* (required for melanin deposition) and tyrosinase (necessary for melanin synthesis) (Julien et al., 2007; Oetting and King, 1999). Melanin loss in the RnPE may have had an additional accelerating role in the degeneration of other eye tissues by leaving them prone to bleaching and oxidative stress which may explain the small and sparse photoreceptors of hagfish (Lamb, 2013). In models of albinism, eye function is seriously impacted in connection with an absence of melanin (Schraermeyer et al., 2006).

Regression of visual cycle machinery at the RPE could also account for the simple complement and small size of hagfish photoreceptors-for example IRBP was the only retinoid cycling constituent not found in the hagfish transcriptome, though it remains to be determined if it is lost from the genome. IRBP is a conserved carrier protein that transports retinoids between photoreceptors and RPE and its loss in mice IRBP slows, but does not eliminate, the recovery of photosensitivity after photobleaching, suggesting that it may not be necessary for retinoid cycling, though it does facilitate a faster restoration of photoreceptor sensitivity (Palczewski et al., 1999; Ripps et al., 2000). Interestingly, IRBP^-/-^ mice have shortened photoreceptors (Ripps et al., 2000), a feature also seen in hagfish.

### Conclusions and limitations

We document that at least one species of extant Hagfish possesses several previously hidden eye synapomorphies; this clarifies the evolutionary origins of the vertebrate eye. Thus, the last common ancestor of agnathans and gnathostomes is inferred to have exhibited the following key characters for vertebrate expansion into diverse photic niches: i) continued retinal growth late into ontogeny supported by a stem cell niche at the retinal margin; ii) a pigmented RPE supporting a forest of ciliary visual photoreceptors; iii) a complex retinal wiring able to compute and decant the photoreceptor outputs into a representation of the visual scene.

As we do not yet have access to embryos of this species, we cannot know if changes to development are responsible for the absence of lens, pigment, and/or cone photoreceptors in adults. It is additionally possible that regression over evolutionary time has eliminated these structures altogether. Other species of hagfish may give us greater insight into the evolutionary history of degeneration in this lineage. For example, the Atlantic hagfish which belongs to a different genus (*Myxine*), is in some ways even more rudimentary than that of the *E. stoutii* and could provide additional perspective on character absence (Fernholm and Holmberg, 1975). Further work is warranted in appreciating the evolutionary and developmental history of visual photoreceptors of hagfish: particularly as they stand in contrast to the cones and cone-like rods found in several species of lamprey (Dickson and Graves, 1979; Morshedian and Fain, 2015).

The results of this study reiterate the great potential for photoreceptive function in hagfish retinas and strengthen the argument for a common origin of vertebrate eyes from a single ancestor that predated the divergence of the agnathans from the gnathostomes. The rudimentary appearance of the hagfish retina has masked many vertebrate features that are now identified including interneurons, photoreceptors, a CMZ-like region, and an RnPE that contains machinery necessary for retina visual function.

A greater understanding of hagfish eye biology can supplement the findings in lamprey eye biology and better inform interpretations of the eye in the last common ancestor of extant agnathans and jawless vertebrates. In light of our confirmation of the presence of interneurons and a retinal (non)pigment epithelium, the eyes of hagfish appear less far removed from that of other vertebrates. Our findings shift the view of hagfish eye evolution from rudimentary to that of a more sophisticated vertebrate-style camera eye.

## Acknowledgements

We thank Dr. Greg Goss, Dr. Alex Clifford, and Dr. Alyssa Weinrauch for their generosity in sharing precious hagfish tissue. Tom Lisney, Eric Clelland, Kelley Bartlett and the staff at Bamfield Marine Science Centre were critical to facilitating hagfish collection and harvesting, particularly. Thank you to Neel Doshi and Arlene Oatway for her help with microscopy. We appreciate the efforts of Nicole Noel, Spencer Balay, and Michèle DuVal in reviewing early versions of this manuscript. This work was supported by an NSERC Discovery Grants and a Bamfield Marine Science Centre research grant awarded to WTA, and an NSERC CGSM Scholarship awarded to EMD.

## Methods

### Animal ethics and tissue collection

This study was conducted under the approval of the Animal Care and Use Committee: BioSciences (University of Alberta Institutional Animal Care and Use Committee, operating under the Canadian Council on Animal Care), Animal Use Protocol Number: AUP00000077. All animals were collected under a permit issued by the Department of Fisheries and Oceans Canada (XR 59 2017), and under animal use protocol approved by the Bamfield Marine Sciences Centre (RS-17-14).

Pacific hagfish (*Eptatretus stoutii*) were collected (N=100) from Barkley Sound, Vancouver Island, BC Canada via a bottom-dwelling trap baited with hake. Hagfish were size selected on board the vessel (MV Alta) and brought ashore to Bamfield Marine Sciences Centre (BMSC) to be housed in aerated, outdoor holding tanks receiving flow-through seawater at 12°C, where they held unfed for 1-3 days before tissue collection. Eyes were obtained from animals euthanized by immersion in a lethal dose of fish anaesthetic MS222 (tricaine methanesulfonate) at 4g/L. Eye tissue was also harvested from fish obtained and used by Dr. Greg Goss and his lab members at the University of Alberta in Edmonton.

Eyes collected for RNA extraction were either placed into RNAlater (Invitrogen, Cat. No. AM7020) at 4°C for 1-14 days before storage at −80°C; or flash frozen in liquid nitrogen before storage at −80°C. Eyes used in transmission electron microscopy (TEM) were fixed in a solution of 2.5% glutaraldehyde/2% PFA/0.1M phosphate buffer. Eyes used for *in situ* hybridization or immunohistochemistry were fixed in 4% paraformaldehyde(PFA)/5% sucrose/0.1M phosphate buffer (referred to hereafter as 4% PFA).

### Tissue preparation for cryosectioning

Following fixation in 4% PFA, tissue was cryopreserved by four sequential sucrose washes: 1 each in 12.5% sucrose/0.1M phosphate buffer and 20% sucrose/0.1M phosphate buffer, overnight in 30% sucrose/0.1M phosphate buffer, and then 1 hour in 1:1 ratio of 30% sucrose/0.1M phosphate buffer:OCT (VWR, Cat No. 25608-930). Tissue was embedded inside a mold made by the severed end of a 1.5mL microcentrifuge tube attached to a glass slide with nail polish and stored at −80°C until used. Tissue blocks were brought to −20°C, sectioned at 10μm at −20°C on a Leica CM1900 UV cryostat then mounted onto SuperFrost Plus glass slides (Fisher Scientific, Cat No. 12-550-15). Sectioned slides were air dried at room temperature and stored at −80°C until use.

### RNA extraction

RNA was extracted using TRIzol (Invitrogen, Oregon, Cat. No.15596026) chloroform extraction procedure (750μl of TRIzol per 50mg of tissue, 0.2mL of chloroform per 1mL TRIzol). Tissues were homogenized using POLYTRON PT 1200 hand held homogenizer (Kinematica Inc.). Chloroform phase separation was followed by centrifugation at 12 000xg for 15 min and then washed with 75% EtOH. RNA quality was assayed using Agilent 2100 Bioanalyzer (Agilent RNA 6000 NanoChip), with RNA Ingegrity Numbers (RIN) of 7 or higher used for downstream experiments.

### Riboprobe production and in situ hybridization

Our *In situ* hybridization is based on a previously described protocol (Barthel and Raymond, 2000). Riboprobes were made by PCR product (cDNA template - qScript cDNA Supermix (Quanta Biosciences, Cat. No. 95048-025) amplified with probes designed against transcripts identified from RNA-sequencing and used to amplify template from cDNA (Key Resources Table). Reverse transcription of PCR products with either DIG or FLR labelled nucleotides by T7 occurred at 37°C overnight. RNA product was precipitated by ethanol at −80°C and final DIG- or FLR-labelled riboprobe quality was assessed by RNA gel electrophoresis on a 1% gel and final concentration was determined by Nanodrop spectrophotometry (GE Healthcare, Cat. No. 28 9244-02).

For *in situ* hybridization, sections were brought to room temperature after storage at −80°C and then fixed for 10 min in 4% PFA at room temperature (RT) followed by proteinase K digestion for 5 minutes, and a second fixation in 4% PFA for 10 minutes at RT. Sections were incubated with 0.3% acetic anhydride and 0.1M triethanolamine for 10 min at, then dehydrated by an ethanol series (50%, 75%, 90%, 100%) and air dried at RT. Hybridization occurred at 65-74°C (probe dependent) for 12-20hrs. Sections were then washed at 65-72°C in 2xSSC 5 min, 0.2xSSC + 0.1% Tween-20 for 20 min, 0.2xSSC + 0.1% Tween-20 2×20 min.

After blocking for at least 60 minutes at RT using 10% normal goat serum in phosphate buffered saline, pH 7.4 (PBS) with 1% Tween-20, sections were incubated in a 1:1000 dilution of alkaline phosphatase-conjugated anti-DIG antibody (Roche Diagnostics, Cat. No. 11207733910) then stained with nitro blue tetrazolium (Roche Diagnostics, Cat. No. 11383213001) and 5-bromo4-chloro-3-indolyl phosphate toludinium (Roche Diagnostics, Cat. No 11383221001). Images were taken on a Zeiss Axio Compound Light Microscope with Optronics MacroFire Digital Camera.

For fluorescent *in situ hybridization*, tissue was incubated with POD (horseradish peroxidase) conjugated anti-DIG or anti-FLR secondary antibodies (1:100 dilution, Roche Diagnostics, Germany, Cat. No. 11093274910, 11426346910) at 4 °C overnight. The following day, tissue was incubated in 1:100 tyramide conjugated to AlexaFluor 488, or 555 (Thermo Fisher Scientific, Cat. No. B40953, B40955). Following development with POD conjugated antibody POD was deactivated by incubation in 3% H_2_O_2_ in PBSTw for 30 minutes at room temperature. Slides were coverslipped with 70% glycerol and sealed with nail polish for imaging.

### Immunohistochemistry

Fronzen tissue sections were brought to RT, and fixed with 4% PFA for 10 minutes, then rinsed with PBSTw. Primary antibodies (Key Resources Table) were applied to tissue and incubated overnight at 4°C and then rinsed with PBS/0.1%Tween 3x 15 mins. Following that, secondary antibody was applied and incubated overnight at 4°C and then rinsed off with PBS/0.1%Tween 3x 15 mins. Secondary antibody used was either AlexaFluor-647 anti-mouse-IgG (Invitrogen, Cat. No. A31571) or AlexaFluor-488 anti-mouse-IgG (Invitrogen, Cat. No. A21202). Secondary antibodies were applied at a dilution of 1:1000 prepared in PBS/0.1% Tween. Sections were then coverslipped with 70% glycerol and sealed with nail polish. To assure specificity, negative controls were performed where no primary antibody omitted.

### Hematoxylin and eosin staining

Tissues were fixed in 4% PFA and embedded in paraffin wax. Fifteen μm sections were mounted onto glass slides. Wax was removed by rinses with toluene. Hematoxylin Gill III (Surgipath, Leica) was applied to the tissue for 2 minutes, followed by 15-minute rinse in distilled water and 2 minutes in 70% ethanol. Eosin (Surgipath, Leica) was applied for 30 seconds, followed by 100% ethanol rinses and 2x 2min toluene rinses. Slides were coverslipped with DPX (Thermo Fisher Scientific).

### Transmission electron microscopy

Tissue preparation and thin sectioning for TEM was performed at the University of Alberta Advanced Microscopy Facility with the help of Arlene Oatway.

After fixation, tissues were stained with 1% osmium tetraoxide/0.1M phosphate buffer at room temperature. Eyes were then embedded in Spurrr’s resin and sectioned at 70-90nm thickness. Sections were further treated with uranyl acetate and lead citrate at room temperature. Micrographs were taken with a Philips-FEI Transmission Electron Microscope (Morgagni 268, operating at 80kV, Hillsboro, Oregon) with Gatan Orius CCD Camera.

### RNAseq

#### Transcript isolation and sequencing

Enucleated eyes were isolated from surrounding muscle in the head. Extraneous tissue was trimmed away. Entire eyes (retina, and vitreous surrounded by a substantial network of collagen and fat) were pooled into a single homogenate and sent for further tissue processing and sequencing procedures at the BGI (Beijing Genomics Institute) at the Children’s Hospital of Philadelphia (CHOP) Genome Center.

#### Methods carried out at the BGI@CHOP Genome Center

Total RNA was extracted with TRIzol (Invitrogen, Oregon, Cat. No.15596026), and quality checked using Agilent 2100 Bioanalyzer (Agilent Technologies, CA, USA). cDNA was synthesized from samples with RIN values of greater than 7. Fragments were sequenced using Illumina HiSeq 4000.

Low quality, adaptor polluted reads, and reads composed of more than 10% unknown bases were filtered by BGI internal software and eliminated from further analysis. Trinity software (v2.0.6) (Grabherr et al., 2011) was used to perform de novo assembly with clean reads. Next, TGicl software (v2.0.6) was used to cluster transcripts and form unigenes (Pertea et al., 2003)

BLAST (v2.2.23) was used to annotate Unigenes by aligning sequences to NT, NR, COG, KEGG and SwissProt databases. Blast2GO (v2.5.0) was used to obtain GO (gene ontology) annotation, and InterProScan5 (v5.11-51.0) was used to attain InterPro annotation {Altschul:1990dw, Altschul:1997td, Conesa:2005hq, Jones:2014fn}. The segment of Unigene that best mapped to functional databases (in order of priority: NR, SwissProt, KEGG, COG) was selected as its coding sequence (CDS). CDS for Unigenes that could not be aligned to any of these databases was predicted by ESTScan (v3.0.2), with Blast-predicted CDS as a model (Christian Iseli, 2002). To determine gene expression, clean reads were mapped to Unigenes using Bowtie2 (v2.2.5), and FPKM gene expression levels were calculated with RSEM (v1.2.12) (Langmead and Salzberg, 2012; Li and Dewey, 2011).

### Sequence homology searching, transcript alignment and tree making

Following *de novo* transcriptome assembly, transcript annotation with 7 functional databases was performed (NR, NT, GO, COG, KEGG, Swissprot, Interpro). To confirm annotations of transcripts of interest, known protein coding sequences from a representative cross section of vertebrates (including zebrafish, mouse, chicken, and frog) were used to search the hagfish eye transcriptome using BLAST 2.6.0 (Altschul et al., 1997) and HMMER 3.1b1 (Eddy, 2011). The top three highly matched sequences were retained, and aligned to homologs using MAFFT under default settings (Katoh et al., 2002; Pearson, 2013). For genes that are part of large and closely related protein families, transcript identity was further confirmed by phylogenetic analysis conducted using IQ-TREE (Nguyen et al., 2015). Branch support was determined using ultrafast bootstrap (1000 bootstrap replicates) (Hoang et al., 2018; Nguyen et al., 2015). The top candidate was used in subsequent analyses.

### Microscopy

Fluorescent micrographs were taken using an LSM 700 inverted confocal microscope in combination with Zeiss Axio Observer.Z1, and captured using ZEN 2010 software (version 6.0, Carl Zeiss AG, Oberkochen). Images were compiled and constructed in PowerPoint (version 1710, Microsoft Office 365). Brightness of images was increased in ImageJ (Schneider et al., 2012).

## KEY RESOURCES TABLE

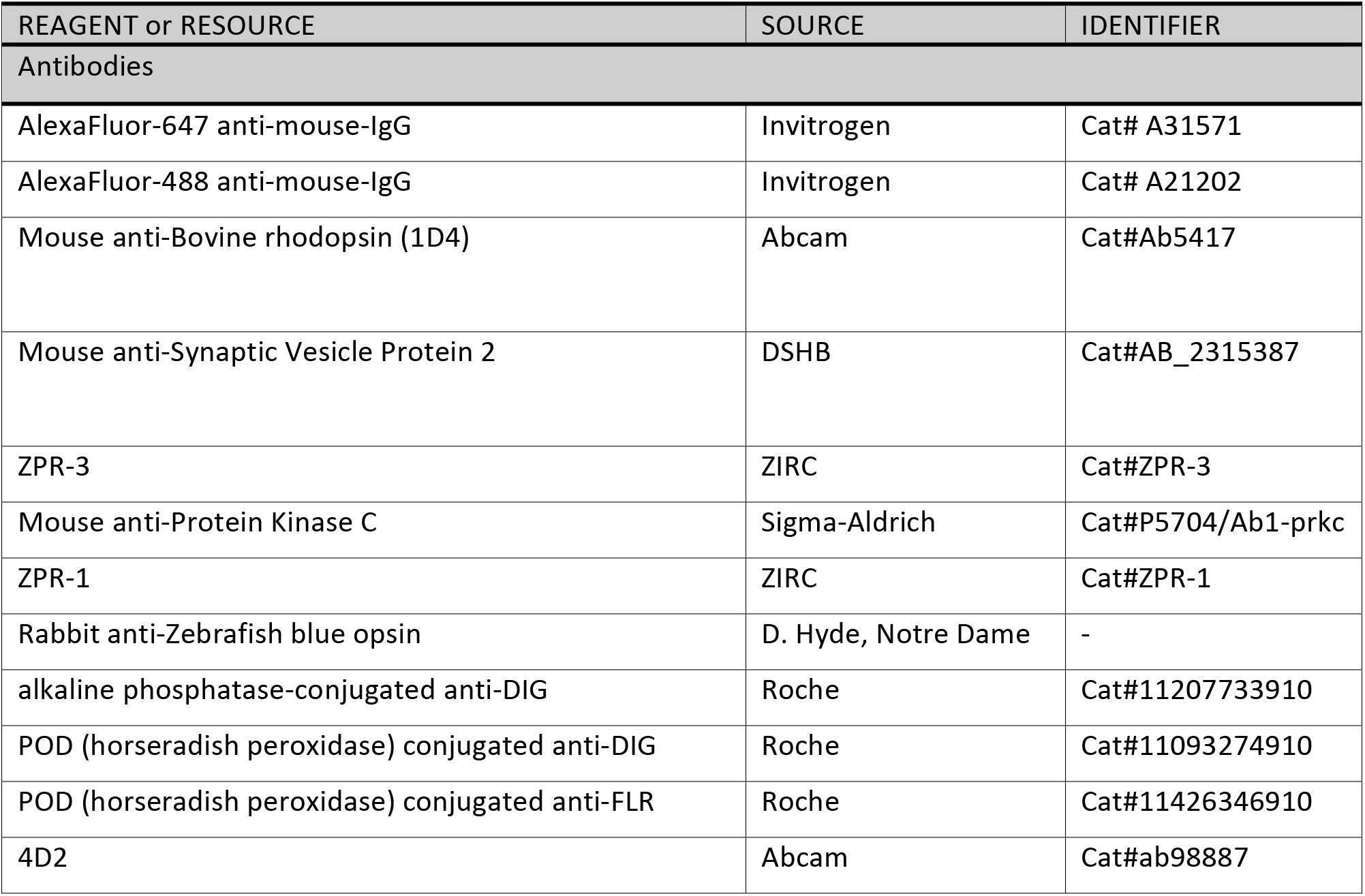
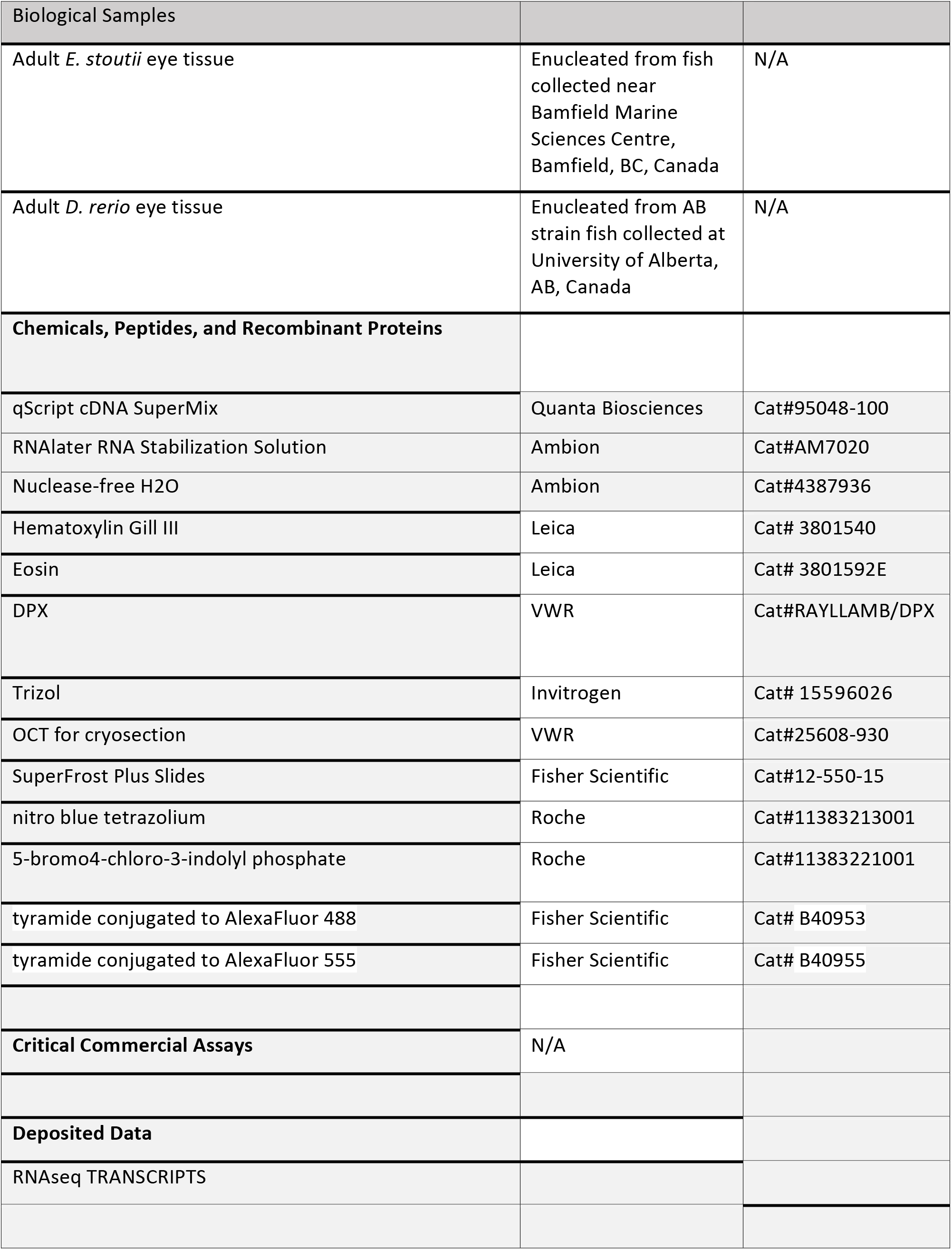
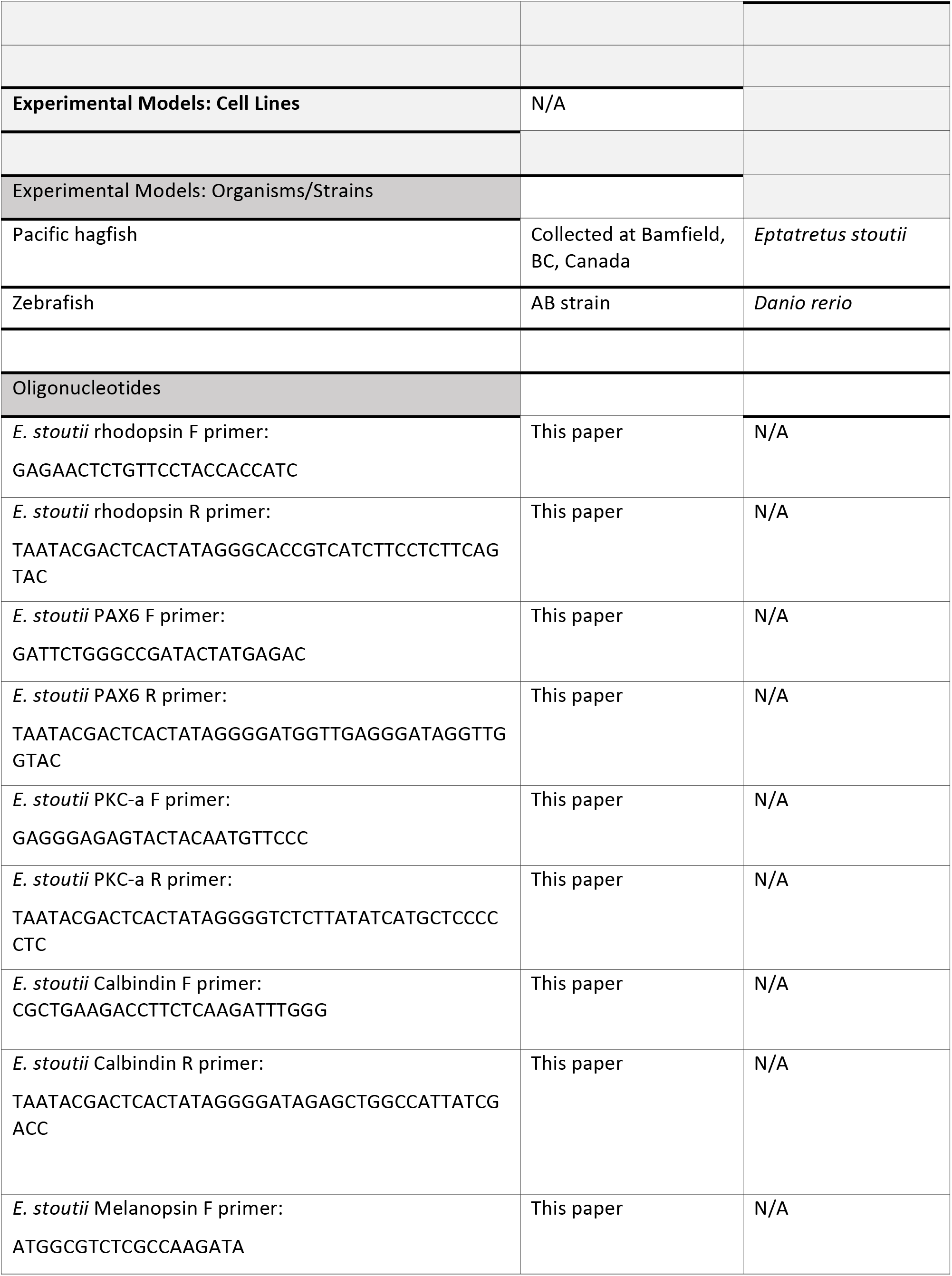
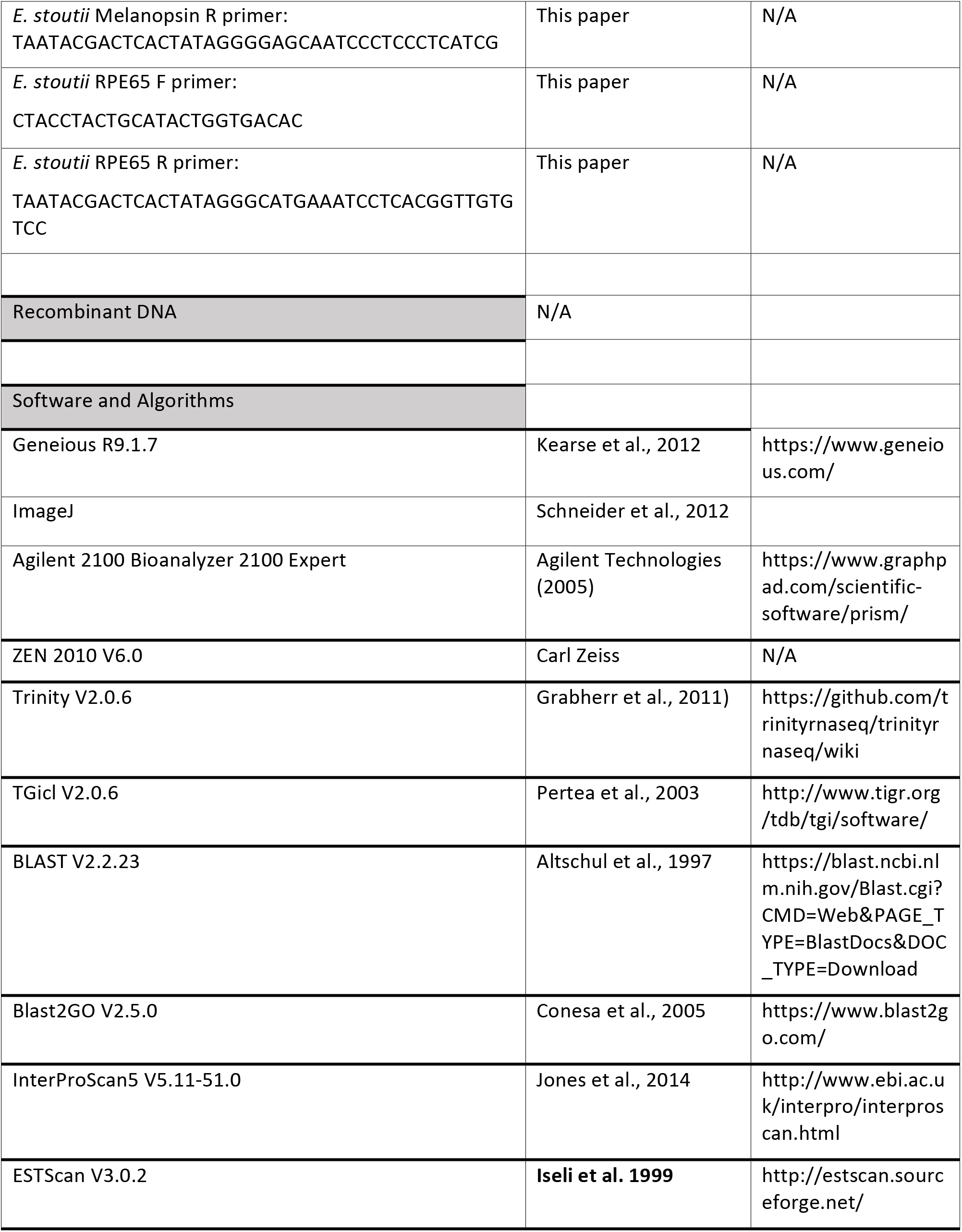
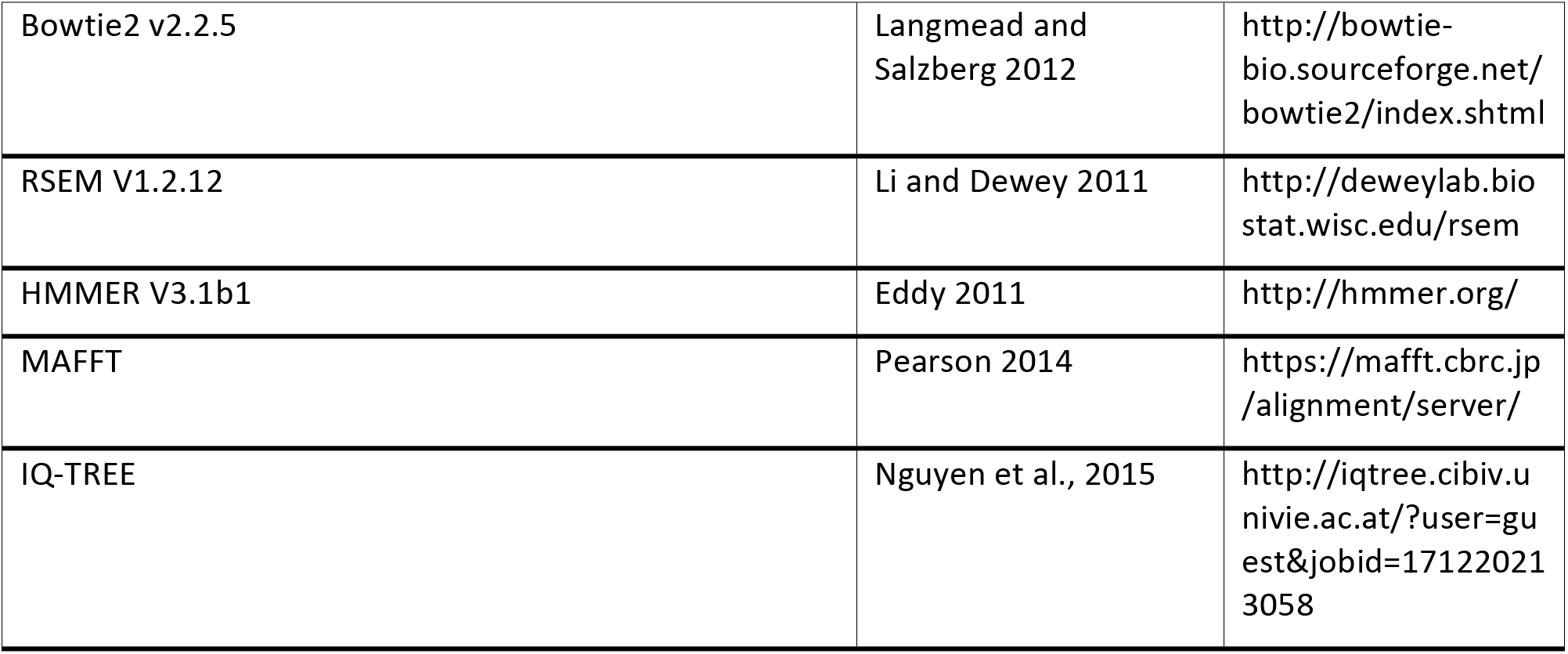

## Supplement to

**Supplemental Figure 1.**
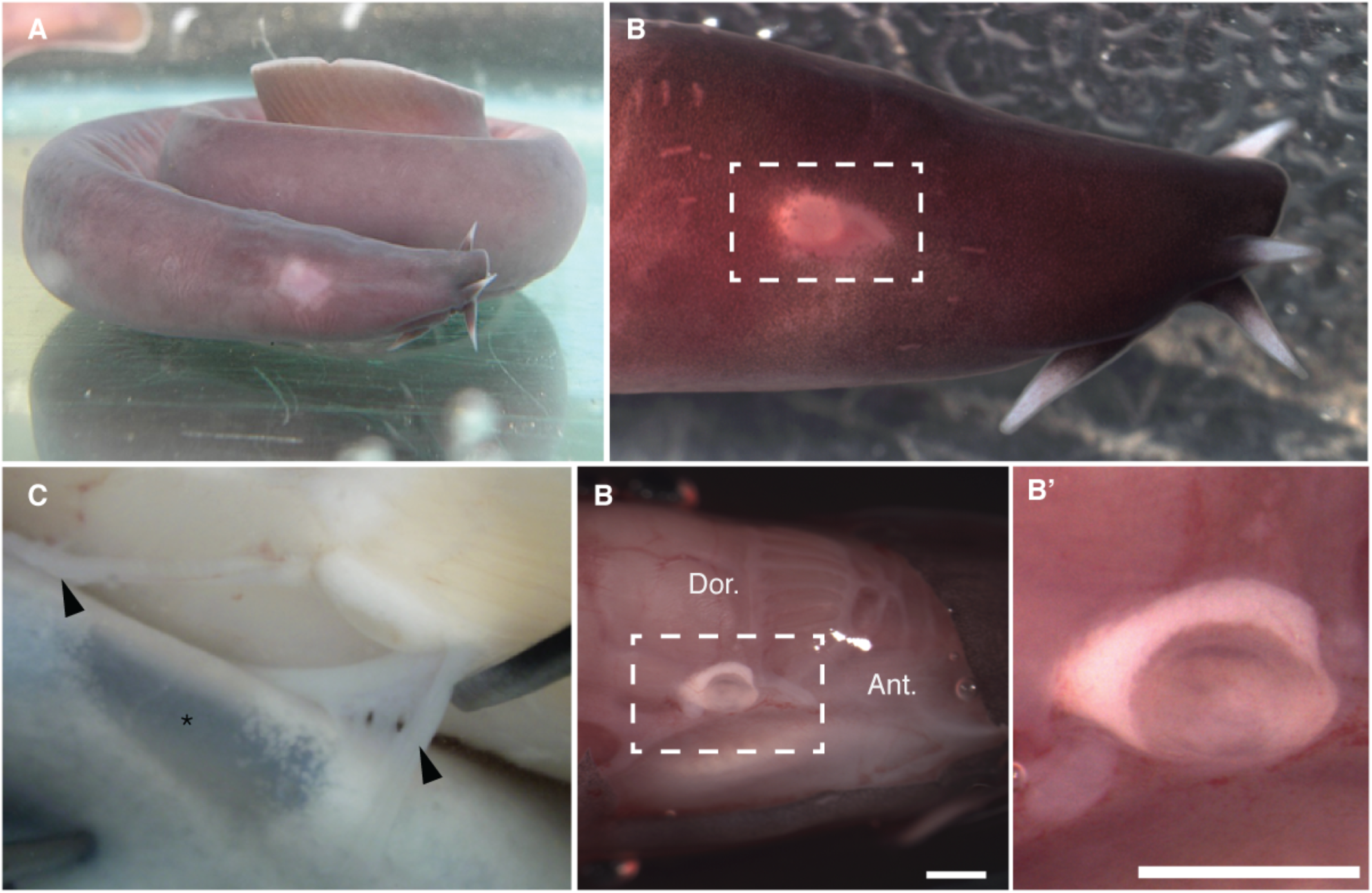
Eye of pacific hagfish is starkly white due to absence of pigmentation. Accompanies Figure 1. **(A)** Live (anaesthetized, intact) hagfish head in lateral view with anterior/snout to the right. Posterior to the tentacles on the snout is a translucent eye patch, and the white of the eye is apparent at the top of this translucent window, indicated by the white box. **(B)** A closer lateral view of the eye of the same hagfish after dissection to remove the skin. B’ is a closer view of the eye.

**Supplemental Figure 2.**
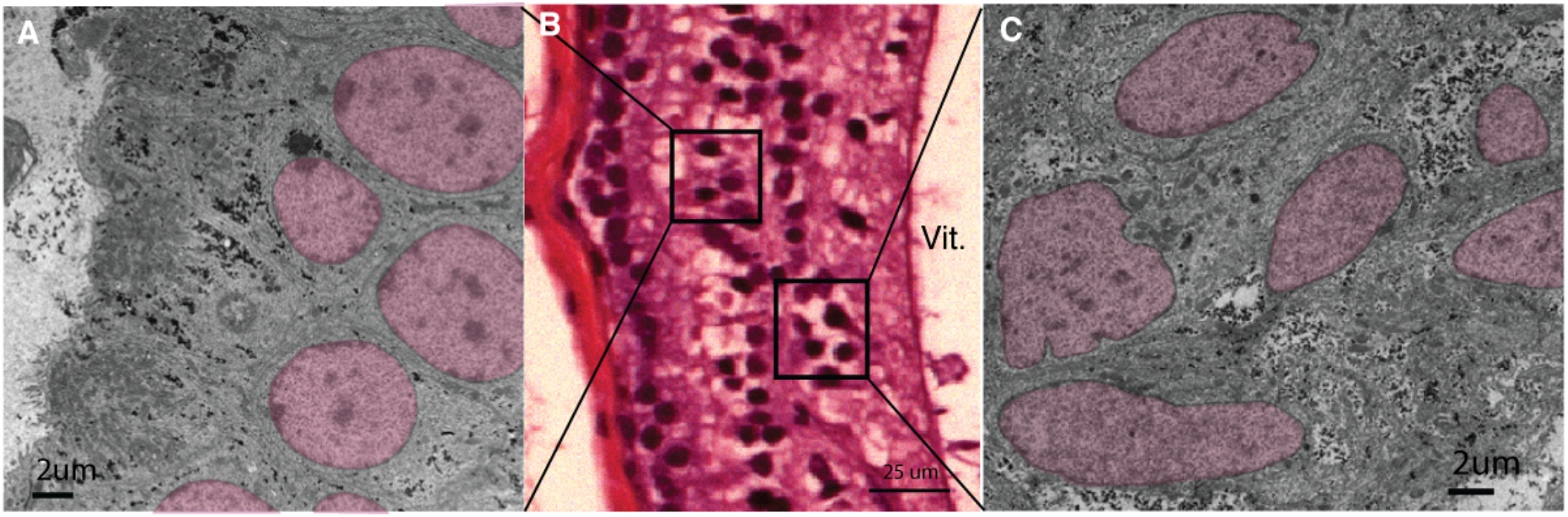
Inner retinal nuclei of hagfish retina are a non-homogenous group of cells. Accompanies Figure 3. Transmission electron microscopy, and H&E stain light microscopy of hagfish retinal nuclei. Photoreceptor nuclei from the photoreceptor layer are consistently ovular shaped and are similarly sized. The cells found vitreal to the photoreceptors are a variety of nuclei that differ in shape and size, an indication that the cells within the inner layer perhaps are not a homogenous population.

**Supplemental Figure 3.**
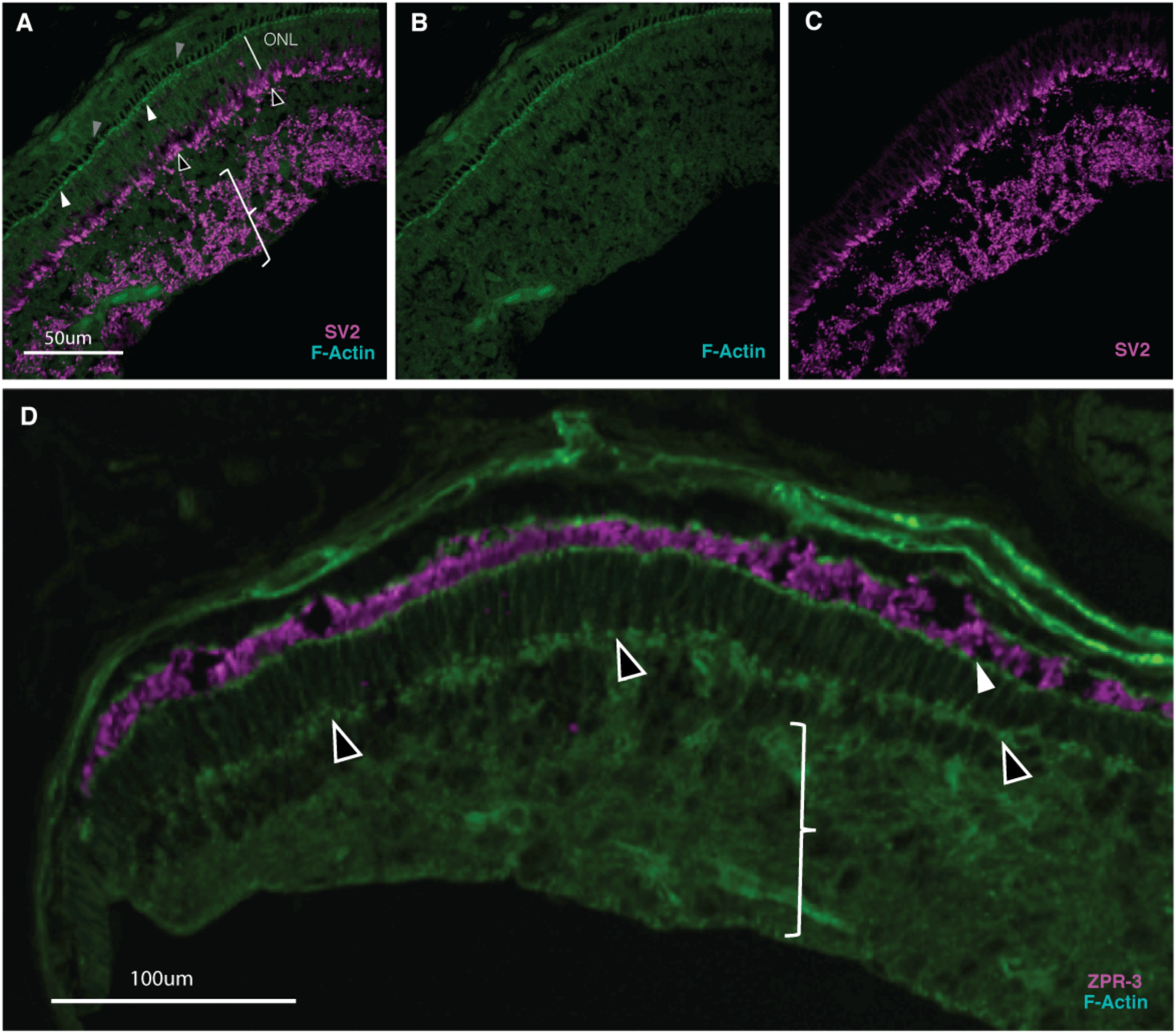
Pacific hagfish neural retina has two plexiform layers. Accompanies Figure 3. **A-C**. Tissue immunoreactive for SV2 (Synaptic vesicle glycoprotein 2A; magenta) indicates synaptic sites. One layer of synapses (empty arrowheads) appears immediately basal of the photoreceptor cells in the outer nuclear layer (ONL). A second layer of labelling occurs at the basal extreme (vitreal aspect) of the retina, amongst the inner retina nuclei (indicated by bracket). Tissue is counterstained with fluorescent phalloidin to label F-actin; F-actin is abundant in a line through the photoreceptors (solid white arrows) and is presumably an equivalent to the Outer Limiting Membrane (OLM) in gnathostome retina, which is composed of Müller glia endfeet. Apical portions of the photoreceptors can be observed immediately apical to and contiguous with the OLM, they appear as a row of gaps in the phalloidin staining (grey arrows in panel A, or are immunostained (magenta) by anti-rod-opsin ZPR-3 (panel **D**). **D**. F-actin is enriched in synapses and various sites of typical nervous system, and in hagfish retina supports the existence of two synaptic plexiform layers (empty arrows and bracket equivalent to panel A).

**Supplemental Figure 4.**
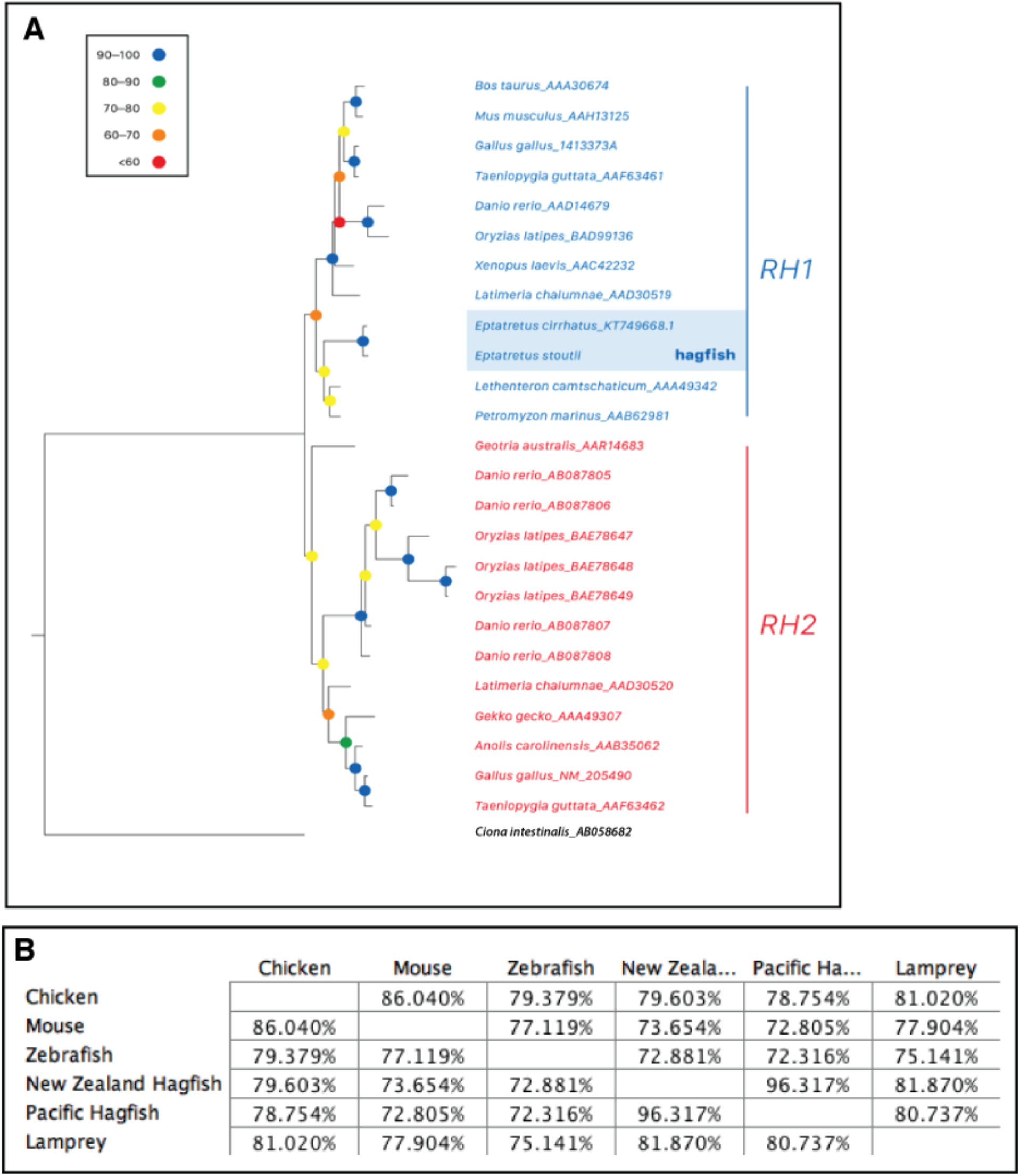
Hagfish rhodopsin is a vertebrate rhodopsin. Accompanies Figure 4. (A) Maximum likelihood tree using protein sequences, including Pacific hagfish rhodopsin, highlighted in blue. (B) Distance matrix (% identity) for protein alignment in B. Node strength is indicated by ultrafast bootstrap support (%).

**Supplemental Table 1.**
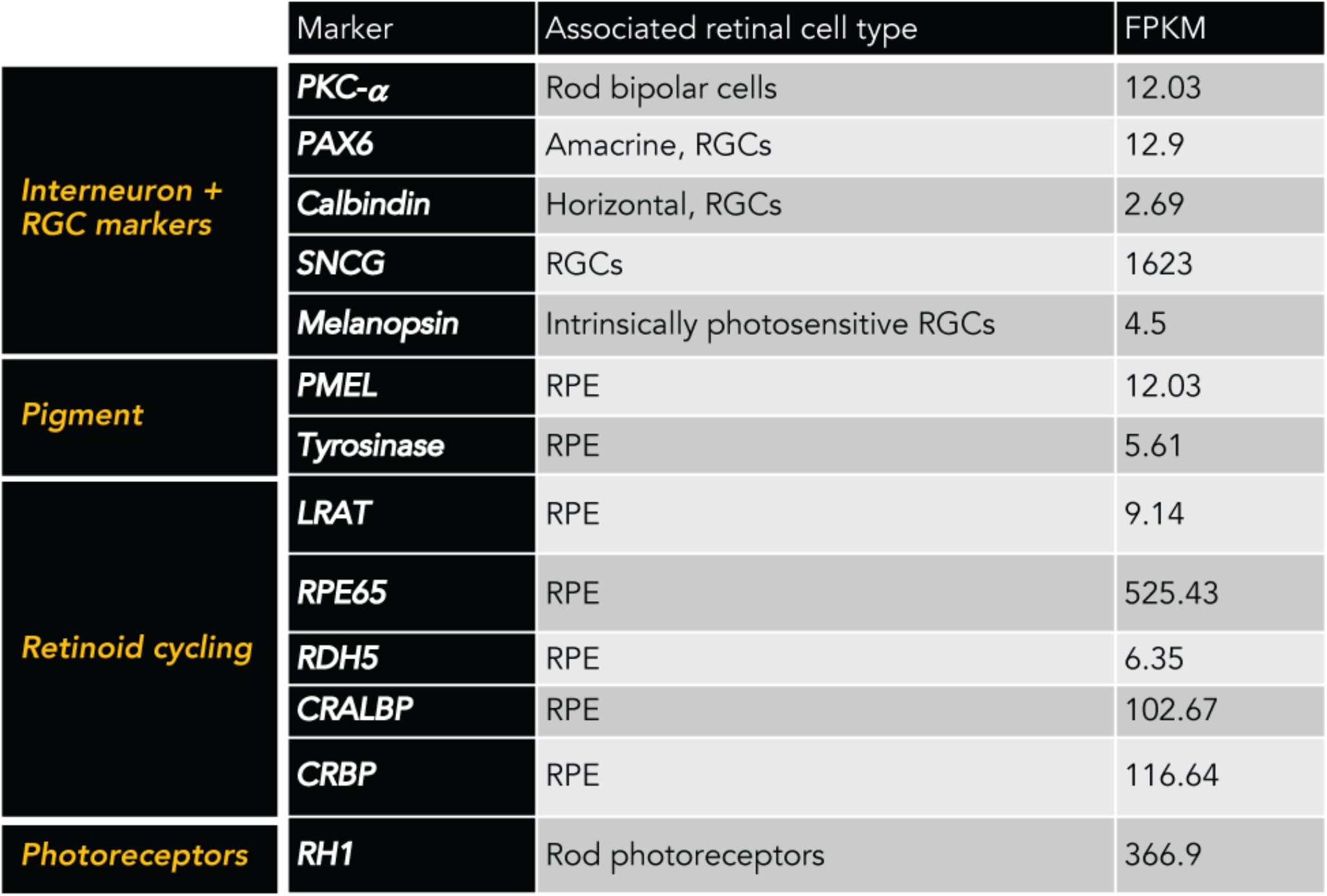
Markers of retinal types found in hagfish

|  | Marker | Associated retinal cell type | FPKM |
| --- | --- | --- | --- |
| **Interneuron + RGC markers** | **PKC-α** | Rod bipolar cells | 12.03 |
|  | **PAX6** | Amacrine, RGCs | 12.9 |
|  | **Calbindin** | Horizontal, RGCs | 2.69 |
|  | **SNCG** | RGCs | 1623 |
|  | **Melanopsin** | Intrinsically photosensitive RGCs | 4.5 |
| **Pigment** | **PMEL** | RPE | 12.03 |
|  | **Tyrosinase** | RPE | 5.61 |
| **Retinoid cycling** | **LRAT** | RPE | 9.14 |
|  | **RPE65** | RPE | 525.43 |
|  | **RDH5** | RPE | 6.35 |
|  | **CRALBP** | RPE | 102.67 |
|  | **CRBP** | RPE | 116.64 |
| **Photoreceptors** | **RH1** | Rod photoreceptors | 366.9 |

